# Search-software choice shapes the discoverable androgen receptor proximal interactome in DIA proteomics

**DOI:** 10.64898/2026.11.15.766752

**Authors:** Lilliana Radoshevich, Michael E. Wright

## Abstract

FragPipe 24.0 (MSFragger-DIA followed by DIA-NN) and Spectronaut 21 (Pulsar/Kuiper) are leading data-independent acquisition (DIA) platforms. Published comparisons differ in library construction and in whether cross-run transfer features such as Match-Between-Runs or background-signal imputation are enabled, confounding these choices with search-engine performance. Here we compare them under a confounder-controlled design, each searching its own fraction-restricted library with cross-run normalization enabled, on a timsTOF Pro2 dia-PASEF dataset of the androgen receptor (AR) proximal interactome across three subcellular compartments. With cross-run transfer symmetrically disabled, Spectronaut 21 identified 24 to 38% more proteins than FragPipe 24.0 in every fraction, and per-run quantities were correlated (Spearman 0.79 to 0.84). FragPipe 24.0 produced tighter biological-replicate CVs (by 4.1 to 6.4 points), while Spectronaut imputation narrowed this gap. Uniquely recovered proteins were at the low-abundance detection limit, yet the two proteomes were compositionally equivalent to within five percentage points, confining divergence to platform-specific sets. Both recovered most of the 989 known AR-interactome members, but platform-unique interactors numbered 34 for Spectronaut 21 and 3 for FragPipe 24.0, with more AR-signaling literature on the Spectronaut-unique side (545 versus 23 co-citing papers). From identical raw data, the platforms agree on most proteins but diverge into complementary, not redundant, platform-specific sets, so running both maximizes discoverable biology.

## Introduction

Data-independent acquisition (DIA) mass spectrometry has transformed quantitative proteomics over the past decade, offering the reproducibility of targeted approaches and the coverage of shotgun methods (1–4). High-throughput DIA workflows on modern instruments, particularly Bruker timsTOF systems using dia-PASEF acquisition (5, 6) and Thermo Orbitrap instruments (7), now routinely quantify thousands of proteins per run across large cohorts, making DIA the platform of choice for both academic biological discovery and industrial biomarker development (8, 9). The bottleneck has shifted from data acquisition to data analysis. How researchers search, score, and integrate raw MS/MS spectra across runs determines what biology is ultimately discoverable. Because different analysis platforms can turn the same spectra into different protein lists, search software is the linchpin of this process, yet researchers rarely examine its influence on what biology a study can ultimately uncover, even as DIA proteomics increasingly drives biological and biomedical discovery.

Two DIA analysis platforms dominate contemporary practice. FragPipe 24.0 (10) integrates the MSFragger fragment-ion-indexed database search engine with DIA-NN (11) for spectrum-centric, library-based quantification, along with tools such as diaTracer (12), MSBooster (13), Percolator (14), and EasyPQP for rescoring and library generation (15). Spectronaut 21 (Biognosys AG) (7) implements the Pulsar and Kuiper search engines (16). It offers a directDIA+ mode that builds a project-specific spectral library from the DIA data itself before targeted extraction, distinguishing it from strictly library-based approaches. Both platforms have matured rapidly, and many laboratories now use them interchangeably, although the choice between them is far from arbitrary. Each uses different algorithms for spectrum-to-peptide matching, different strategies for cross-run transfer of identifications, and different defaults for FDR estimation (17), all of which can shape the final protein list in ways not immediately transparent to the end user.

Several recent studies have compared these platforms head-to-head (18–21). Zhang et al. (22) benchmarked five DIA analysis tools (OpenSWATH (23), EncyclopeDIA (24), Skyline (25), DIA-NN, and Spectronaut) across six datasets acquired on TripleTOF, Orbitrap, and timsTOF Pro2 instruments, comparing precursor and protein identifications and quantification consistency. Their central methodological finding was that library-free approaches outperform library-based ones (26) when the reference spectral library is substantially smaller than the sequence space. Yu et al. (15) introduced MSFragger-DIA and validated it against Spectronaut and DIA-NN on multiple datasets, reporting comparable identification depth, tighter FDR control, and shorter runtimes. Demichev et al. (27) established the dia-PASEF workflow in FragPipe and DIA-NN for timsTOF data, achieving up to 9000 quantified protein groups per run on HeLa lysates. More recently, Koudelka et al. (28) directly compared FragPipe 21 (MSFragger 4.0 → DIA-NN 1.8.2) with Spectronaut 18 in the context of limited proteolysis mass spectrometry (LiP-MS) for drug-target identification, concluding that “the choice between precision (FragPipe) and sensitivity (Spectronaut) largely depends on the specific experimental context.” Two recent reviews (8, 9) later synthesized this emerging consensus and framed platform choice as an open question, one that depends on both the experimental aim and downstream analytical goals.

Despite their collective value, three gaps remain. First, all published head-to-head comparisons, including those cited above, have run each platform at its default configuration. In DIA-NN, match-between-runs (MBR) is a distinct, toggleable step that transfers identifications between runs (11). It is commonly enabled when DIA-NN is run on its own, although the DIA-NN packaged within FragPipe leaves it off by default. Spectronaut’s directDIA performs cross-run identification as an intrinsic part of its analysis rather than as a separate rescue and does not apply an MBR-style transfer by default, so its closest analog to MBR is the optional background-signal imputation applied at the quantification stage (7). Because a cross-run rescue can add identifications or fill quantities not independently confirmed by each run’s MS/MS evidence, any observed difference in platform depth measured at defaults conflates the search engine’s contribution with that of the cross-run strategy. Isolating the algorithmic contribution requires symmetrically controlling cross-run transfer on both platforms, a matched configuration that no published benchmark has systematically applied. Second, all published comparisons operate on a single biological sample or dataset, so observed platform differences cannot be distinguished from dataset-specific idiosyncrasies. The literature lacks comparisons across multiple biological conditions within the same underlying system. Third, and most importantly, prior comparisons focus on counts (precursors identified, proteins quantified, coefficients of variation across replicates) but do not ask whether proteins that differ between platforms represent different biology or the same biology detected at different rates. To our knowledge, no one has systematically characterized the biological content of the divergence zone between platforms.

Here, we addressed these three gaps through a confounder-controlled, head-to-head comparison of FragPipe 24.0 (FP-24.0; MSFragger-DIA + DIA-NN 1.8.2 beta 8) and Spectronaut 21 (SN-21; Pulsar/Kuiper), evaluating protein identification and protein-level quantification on a Bruker timsTOF Pro2 dia-PASEF dataset of the androgen receptor (AR) proximal interactome mapped by APEX2 proximity labeling across an androgen-stimulation time course (29). AR is a ligand-activated transcription factor whose regulatory network includes nuclear cofactors, chromatin modifiers, cytoplasmic scaffolds, and membrane-proximal signaling proteins, the very categories whose recovery in DIA-MS depends most on the analysis platform’s sensitivity. The analysis was replicated across three biochemically distinct subcellular fractions (63 cytosolic, 85 nuclear, and 63 microsomal DIA runs), providing three independent tests of the same comparative claims. To remove library construction as a confounder, each platform searched its own fraction-restricted library built from that fraction’s DDA and DIA runs, so all three fractions were analyzed identically, and any platform differences reflect the search and quantification engines rather than library design. We controlled cross-run transfer symmetrically, first with the feature disabled on both platforms (Match-Between-Runs in DIA-NN, background-signal imputation in Spectronaut 21) and then with it enabled on both. We report both the identification and quantitative behavior of the two platforms and interrogate the divergence-zone proteins against curated interactor references and the AR-specific published literature.

With cross-run transfer disabled on both platforms, Spectronaut 21 held a substantial depth advantage over FragPipe 24.0, ranging from 24% to 38% across the three fractions. Because library construction was matched, that advantage reflects the search and quantification engines rather than library design. FragPipe 24.0, in turn, was the more precise quantifier, with consistently tighter biological-replicate CVs, even as the two platforms agreed closely on the values they assigned to shared proteins (global Spearman 0.79 to 0.84). The platforms thus diverge in what they identify, not in how they quantify what they share. Most consequentially, each platform’s divergence zone is enriched for a different class of AR biology. On the Spectronaut 21 side, these include transcription factors, mitochondrial proteins, and membrane transporters, whereas on the FragPipe 24.0 side they include cytoskeletal and muscle proteins. Even though the two full proteomes are compositionally equivalent, the divergence is confined to the proteins each uniquely detects rather than spread across the proteome. These divergence-zone proteins sit at the low-abundance edge of the dynamic range rather than being noise, and independent PubMed validation confirms that both zones contain published AR-interactors, with far more cumulative literature on the Spectronaut 21 side. The rankings held with cross-run transfer enabled, where Spectronaut imputation narrowed the CV gap. Running both engines rather than choosing one therefore maximizes recoverable biology from a proximity-labeling DIA experiment, especially from the proximal androgen-receptor interactome.

## Experimental Section

### Experimental Design and Statistical Rationale

This paired platform comparison used identical raw input files for both analysis platforms, so any output differences are attributable to the analysis pipeline rather than the sample or acquisition. We independently analyzed three biochemically distinct subcellular fractions (*i.e.*, cytosolic, nuclear, and microsomal), providing three tests of the same comparative claims. Each fraction comprised seven proximity-labeling conditions along an androgen stimulation time course (a no-hydrogen-peroxide negative control, then 0, 2, 10, 30, 60, and 360 minutes of synthetic androgen R1881, with APEX2 biotinylation triggered by hydrogen peroxide at each labeled time point) in three biological replicates, each sample separated into three high-pH reversed-phase peptide fractions, giving 63 cytosolic (seven conditions x three biological replicates x three peptide fractions), 85 nuclear, and 63 microsomal dia-PASEF runs. The nuclear fraction includes additional biological-replicate reruns.

In the primary comparison, both platforms were configured to a matched baseline in which the platform-specific cross-run transfer feature was disabled (Match-Between-Runs in DIA-NN and no background-signal imputation in SN-21), so that identifications in any given run reflect only that run’s evidence and no imputation was applied. We controlled protein-group identifications at a 1% false discovery rate on both platforms. We analyzed a matched cross-run-transfer-enabled state (Match-Between-Runs in DIA-NN and background-signal imputation in SN-21) in parallel as a robustness check. We chose statistical treatments to match the data’s distributional properties. We computed coefficients of variation per protein group across biological replicates within each condition using only shared protein groups and reported them as medians per platform per fraction, so quantitative comparison is restricted to proteins detected on both platforms. We assessed cross-platform quantitative agreement using Spearman rank correlation on log_10_ MaxLFQ (30) intensities, a rank-based measure that does not assume normality. We compared intensity and per-run completeness distributions between the shared, SN-only, and FP-only sets using the Mann-Whitney U test. We compared contaminant enrichment against fraction baselines using Fisher’s exact test. We performed functional enrichment using g:Profiler against a fraction-specific custom background comprising the full protein set measured in that fraction, with a Benjamini-Hochberg false discovery rate threshold of 0.05, and we enabled cross-run normalization on both platforms throughout.

### Cell culture and sample preparation

We generated the experimental dataset by quantitatively mass-spectrometrically profiling the androgen receptor (AR) proximal interactome in LNCaP prostate tumor cells that stably express a doxycycline-inducible APEX2-AR fusion protein (31). APEX2-AR expression was induced with 10 ng/mL doxycycline; cells were incubated with 500 µM biotin-tyramide for 1 hour at 37°C, and the synthetic androgen R1881 was added at 100 nM for a time course of 0, 2, 10, 30, 60, and 360 minutes. APEX2-catalyzed proximity biotinylation was triggered by a 60-second pulse of 1 mM hydrogen peroxide; a parallel no-hydrogen-peroxide sample, referred to as ‘minus,’ served as the negative labeling control. Samples were collected in three biological replicates (R1, R2, R3) and fractionated into cytosolic, nuclear, and microsomal subcellular fractions. For each time point, approximately 5 mg of input protein was diluted into denaturing binding buffer (100 mM EPPS, 400 mM NaCl, 4 M urea, 1% SDS, pH 8.5), reduced with 5 mM TCEP, alkylated with 10 mM chloroacetamide, and quenched with 5 mM DTT. Biotinylated proteins were then captured on Pierce magnetic streptavidin beads, and the carbamidomethylated, trypsin-digested peptides were desalted, fractionated by offline high-pH reversed-phase chromatography into three peptide fractions per sample, aliquoted, and stored at -80°C until acquisition, with a Pierce retention time calibration (PRTC) peptide standard spiked into each sample for retention-time calibration.

### dia-PASEF LC-MS/MS acquisition

Samples were acquired on a Bruker timsTOF Pro2 mass spectrometer coupled to a nanoElute 2 UHPLC in data-independent acquisition parallel-accumulation-serial-fragmentation (dia-PASEF) mode. Isolation windows followed a py_diAID-optimized dia-PASEF scheme (12 to 25 scan windows) over a precursor m/z range of 200 to 1450 and an ion-mobility (1/K0) range of 0.70 to 1.40 V·s/cm², with a 100 ms ramp time and a 100% duty cycle. Chromatographic separation used a 60-minute gradient at 500 nL/min. The complete acquisition schedule comprised 63 cytosolic DIA runs (7 conditions x 3 biological replicates x 3 high-pH reversed-phase peptide fractions), 85 nuclear DIA runs (with additional biological-replicate reruns), and 63 microsomal DIA runs. Both analysis pipelines used raw .d files directly; no format conversion was required.

### Protein sequence database

Both pipelines searched the human UniProt proteome (32) without isoforms (database release 2023-10-10). For FP-24.0, we appended reversed decoy sequences to the target database with the decoy prefix rev_. For SN-21, the forward-only target FASTA was supplied, and SN-21 generated its own decoys internally during the Pulsar search using its swap-after-reverse rule. Both analyses used the same target sequences; decoy handling differed between platforms, as reported in Supplementary Table S2.

### FragPipe 24.0 analysis pipeline

Analyses were performed in FP-24.0 following the MSFragger-DIA hybrid-library workflow described by Yu et al. (15), adapted for timsTOF dia-PASEF data with diaTracer (v2.2.1) performing pseudo-spectrum extraction, the timsTOF-native equivalent of the DIA-Umpire step (33) used on Orbitrap data. The pipeline stages are executed in sequence per fraction. (i) diaTracer generated pseudo-MS/MS spectra from the dia-PASEF .d files using a correlation threshold of 0.3, delta-apex ion mobility of 0.01 (1/K0), delta-apex retention time of 3 scans, and a mass-defect filter with offset 0.1; (ii) MSFragger (v4.4.1) performed a spectrum-centric database search of the diaTracer pseudo-spectra in DDA mode against the decoy-appended human database with a 20 ppm precursor mass tolerance, a 20 ppm fragment mass tolerance, strict trypsin cleavage rules (stricttrypsin enzyme setting) with up to 2 missed cleavages, peptide lengths of 7-50 amino acids, and the modification set described below; (iii) MSBooster (v1.4.14) computed deep-learning features (predicted retention time, fragment-ion spectra, and ion mobility) using the DIA-NN prediction models; (iv) Percolator (v3.7.1) rescored PSMs using the MSBooster features with target-decoy competition; (v) Philosopher (v5.1.3-RC9) (34) performed protein inference (ProteinProphet) (35) and applied picked-protein FDR filtering (36) at 1% at the protein level (--picked --prot 0.01); (vi) EasyPQP (FragPipe-SpecLib v0.1.58) assembled the hybrid spectral library from the combined DDA and DIA-search PSMs, selecting top-6 b/y fragment ions per assay with Pierce iRT reference calibration for retention time and automatic reference-run selection for ion mobility; (vii) DIA-NN (v1.8.2 beta 8) performed targeted extraction and quantification of the primary DIA runs against the EasyPQP hybrid library at a 1% precursor and protein-group q-value threshold, with cross-run normalization set to global default and Match-Between-Runs (MBR) deliberately disabled (diann.mbr = false) as part of the matched configuration described below. Protein-group quantities were taken from the DIA-NN protein-group matrix (report.pg_matrix.tsv) and precursor-level quantities from report.pr_matrix.tsv. In this DIA workflow, DIA-NN performs quantification rather than IonQuant, FP-24.0’s quantifier for data-dependent acquisition, so the reported protein-group and precursor quantities are DIA-NN outputs. The Match-Between-Runs setting in this study therefore refers specifically to DIA-NN’s second-pass match-between-runs (the reanalyse option, controlled by diann.mbr), and IonQuant’s data-dependent-acquisition match-between-runs does not contribute to any reported DIA quantity.

### Spectronaut 21 analysis pipeline

We performed analyses in SN-21 (Biognosys) using the Kuiper search engine, the successor to Pulsar. We analyzed all three subcellular fractions identically using the directDIA+ workflow (Deep setting) and a fraction-restricted hybrid library. For each fraction, that fraction’s DIA runs and matched DDA runs (Supplementary Table S1) were combined into a single one-step analysis. SN-21 built a project-specific hybrid library from that fraction’s DDA and DIA runs and then performed a targeted analysis of the same DIA runs. We enabled the Hybrid (DDA + DIA) Library option for all three fractions. We set source-specific iRT calibration to Auto and disabled the Biognosys iRT Kit setting because the study used the Pierce PRTC rather than the Biognosys kit. Pulsar search settings were matched to the FP-24.0 settings where an equivalent SN-21 parameter existed, namely a closed (non-open) tryptic search with strict trypsin cleavage rules and carbamidomethyl (C) as the fixed modification, with the heavy Pierce PRTC peptides configured through the Labeling tab as a heavy channel (described below). We set FDR to 1% at the PSM, peptide, and protein-group levels; selected MaxLFQ as the roll-up strategy; and enabled cross-run normalization. In the primary comparison, background-signal imputation was disabled, and a matched state with background-signal imputation enabled was analyzed in parallel.

### Modifications applied in both pipelines

We applied a matched set of fixed and variable modifications in both pipelines. The fixed modification was carbamidomethylation of cysteine (+57.021464 Da). The variable modifications were oxidation of methionine (+15.994915 Da, up to 2 per peptide), acetylation of the protein N-terminus (+42.010565 Da, up to 1 per peptide), and phosphorylation of serine, threonine, or tyrosine (+79.966331 Da, up to 3 per peptide), with a maximum of three variable modifications per peptide in both pipelines. We treated the spiked heavy Pierce PRTC peptides as a heavy label rather than as variable modifications. In SN-21, we configured them in the Labeling tab as a second channel carrying heavy Arg10 (+10.008269 Da) and Lys8 (+8.014199 Da), and we in silico generated the missing heavy channel, following Biognosys guidance for a label-free study with a heavy spike-in. In FP-24.0 and DIA-NN, the corresponding heavy Lys8 and Arg10 were declared for identification of the spiked peptides. Phosphorylation was included because AR-interactome biology plausibly involves phosphorylation at S/T/Y sites.

### Matched cross-run-transfer configuration

To separate the contributions of the search-engine and peptide-centric quantification from the platform-specific cross-run transfer feature, we configured the two platforms symmetrically. In the primary comparison, we disabled cross-run transfer on both platforms, with Match-Between-Runs disabled in DIA-NN (diann.mbr = false) and no background-signal imputation in SN-21. In FragPipe 24.0 the packaged DIA-NN runs with Match-Between-Runs off by default, so this primary configuration matches the FragPipe default while removing SN-21’s optional imputation. To characterize the effect of cross-run transfer itself, we also analyzed a matched transfer-enabled state (Match-Between-Runs enabled in DIA-NN, background-signal imputation enabled in SN-21), which is non-default for FragPipe, and we report the comparison at both settings. We enabled cross-run normalization on both platforms throughout. We held all other parameters identical or matched to the closest equivalent between the two platforms per the parameter map (Supplementary Table S2).

### Divergence-zone analysis and functional enrichment

For each fraction, we partitioned protein groups identified at 1% FDR by both platforms into three sets, comprising shared protein groups identified by both platforms, SN-only protein groups identified by SN-21 but not by FP-24.0, and FP-only protein groups identified by FP-24.0 but not by SN-21. We first expanded protein groups reported by each platform to their constituent UniProt accessions because MSFragger with ProteinProphet and Kuiper with IDPicker group shared peptides using different algorithms, so a group string is not comparable between platforms. We report identification counts and divergence zones by accession. For functional enrichment, we mapped accessions to gene symbols and determined set membership at the gene level, so we considered a gene reported by both platforms under different accession counts shared rather than unique to either platform. We performed functional enrichment analysis on each divergence-zone protein set with g:Profiler (37) via the REST API (POST/api/gost/profile/), using a fraction-specific custom background comprising the full protein-group set measured in that fraction. Enrichment sources included GO Biological Process, GO Molecular Function, GO Cellular Component, KEGG, and Reactome. Benjamini-Hochberg FDR < 0.05 was used as the significance threshold, and annotations inferred exclusively from electronic annotation (IEA) were excluded. The full enrichment tables per fraction per set are provided in Supplementary Table S4. Cross-fraction concordance of GO term recurrence and enrichment on recurrent-across-3-fractions gene lists are provided in Supplementary Table S5. Supplementary Table S9 provides enrichment and member-to-member network analyses of interactome members recovered by only one platform.

The whole-proteome comparison between platforms shown in Figure 7 was computed as subcellular compartment composition rather than as differential enrichment. For each fraction, we mapped each platform’s full detected proteome to gene symbols. The fraction of that proteome annotated to each Gene Ontology Cellular Component term was determined with g:Profiler against the annotated human genome (domain scope annotated, electronic annotations excluded), as the number of proteome genes in the term divided by the proteome size. Because this membership measure is independent of the comparator platform and any enrichment background, it reports each proteome’s subcellular composition directly rather than its over-representation relative to a reference set.

To distinguish genuine detection from homology-driven inference for the tissue-restricted proteins in the cytosolic and nuclear FP-only cytoskeletal sets, each gene was checked in the FP-24.0 DIA-NN precursor report (report.pr_matrix.tsv) for a proteotypic precursor, a peptide mapping uniquely to that gene in the search database and quantified in at least one run. We classified a gene with a quantified proteotypic precursor as genuinely detected on unique evidence, and a gene supported only by shared precursors as a homology-driven inference (Supplementary Table S10).

### AR-interactor overlap and PubMed literature validation

We used a reference set of known androgen receptor (AR) interaction partners, a curated 989-member AR-interactome assembled from public protein-protein interaction resources. Because these resources record both physical and genetic interactions, the reference set includes a small number of members whose association with the androgen receptor is genetic rather than physical, for example TP53, which appears through a suppressive genetic interaction. We treat membership uniformly, without stratifying by interaction type. Hence, recovery of a member indicates its presence in the reference set and is not, in every case, evidence of a direct physical interaction. We applied no confidence threshold or abundance filter to the reference set. We mapped protein sets from UniProt accession to gene symbol and intersected them with the reference set. One gene symbol required manual curation. The symbol TCF4 denotes the basic helix-loop-helix factor ITF2 (E2-2), but the androgen receptor literature and reference annotation under that symbol reflect the Wnt effector TCF7L2, which both platforms detected and which is therefore shared rather than platform-unique. We excluded this single artifactual entry from the strict platform-unique network, giving 34 SN-21-unique members rather than 35. We assessed platform membership at the gene level, as in the divergence-zone analysis above. We retrieved PubMed literature co-citation counts via the NCBI Entrez E-utilities API (esearch endpoint) (38) for every interactome member uniquely recovered by one platform, using the query ’[gene symbol] AND androgen receptor’, and froze the results on the query date to ensure reproducibility. Supplementary Table S6 provides per-gene counts and the lists of recovered members.

### Mechanistic characterization of divergence-zone proteins

To assess whether divergence-zone protein groups reflect genuine low-abundance detection or the accumulation of noisy marginal PSMs, we computed three per-protein metrics. (i) median log_10_ protein-group intensity across all quantified runs, used as a per-protein abundance estimate; (ii) per-run completeness, defined as the number of runs (of the 63, 85, or 63 total per fraction) in which each protein group was quantified; and (iii) common-contaminant class membership, determined by matching UniProt accessions against the universal protein contaminant library of 381 entries (39). We compared intensity distributions across shared, SN-only, and FP-only sets per fraction using the Mann-Whitney U test, compared completeness distributions with the same test, and compared contaminant enrichment rates to fraction baselines using Fisher’s exact test. Figure 6 summarizes the results across the three fractions.

### Statistical analysis and reproducibility

We computed Spearman rank correlations of log_10_ MaxLFQ intensities for shared protein groups pairwise across biological replicates and platforms; we report per-run and global values in the Results (Figure 5). We computed the median coefficient of variation per protein group across the three biological replicates using non-imputed intensities. For each biological replicate and hydrogen-peroxide-labeled time point, the protein-group intensities of the three high-pH reversed-phase peptide fractions were summed into a single per-sample value before computing the coefficient of variation, so that reproducibility is assessed at the level of the biological sample rather than the individual peptide fraction. Because we computed MaxLFQ intensities per run rather than jointly across the three fractions, this summation approximates a sample-level quantity. We excluded the no-hydrogen-peroxide negative controls because they contain little labeled protein, and their replicate variation reflects near-baseline signal rather than quantitative reproducibility. Values are reported as medians per subcellular fraction per platform (Figure 4; diagrammed in Supplementary Figure S2). We performed all analyses in Python 3.10 using openpyxl, pandas, numpy, scipy. We used matplotlib, seaborn, and networkx, and called the g:Profiler REST API via urllib.request. We provide software versions and analysis scripts in Supplementary Table S8. Primary mass spectrometry data are available as described in the Data Availability statement (ProteomeXchange PXD081449).

### Spectral library composition

We built spectral libraries strictly on a per-fraction basis, constructing each platform’s library for a given fraction from only that fraction’s DDA and DIA runs, with no cross-fraction DDA or DIA content at any stage. The cytosolic library comprises 105 cytosolic DDA files from three biological replicates and 63 cytosolic DIA runs. The nuclear library comprises 63 nuclear DDA files and 85 nuclear DIA runs. The microsomal library comprises 21 microsomal DDA files and 63 microsomal DIA runs. This fraction-restricted design applies the same library-construction rules to every fraction and both platforms, so platform comparisons are not confounded by library-design differences.

No unexpected, new, or significant hazards or risks were associated with the reported work. Standard laboratory safety practices were followed for mammalian cell culture and for the brief use of hydrogen peroxide during proximity labeling.

## Results

### Under Matched Conditions, the Engines Differ Substantially in Identification Depth

#### Study design and confounder control

FragPipe 24.0 (FP-24.0; MSFragger-DIA plus DIA-NN 1.8.2 beta 8) and Spectronaut 21 (SN-21; Pulsar/Kuiper) are usually compared without matching library construction or symmetrically controlling cross-run transfer, so each search engine stays entangled with those choices. We set out to disentangle them by measuring what each engine recovers on its own, separate from the transfer features. We analyzed a single Bruker timsTOF Pro2 dia-PASEF dataset of the AR proximal interactome (29) across its cytosolic, nuclear, and microsomal fractions (Figure 1), holding every analysis choice constant except the engines. Each platform searched its own fraction-restricted library, built the same way for both and never drawing on another fraction’s runs, and we first disabled every cross-run transfer feature (Match-Between-Runs in DIA-NN, background-signal imputation in SN-21) so that each identification rested on its own run alone. We then repeated the comparison with transfer enabled on both sides. Any difference that survives this design reflects differences in the search and quantification engines rather than library construction or gap-filling, which is the basis on which we compare the proteins each engine identifies and how closely they agree on the proteins both detect.

**Figure 1.**
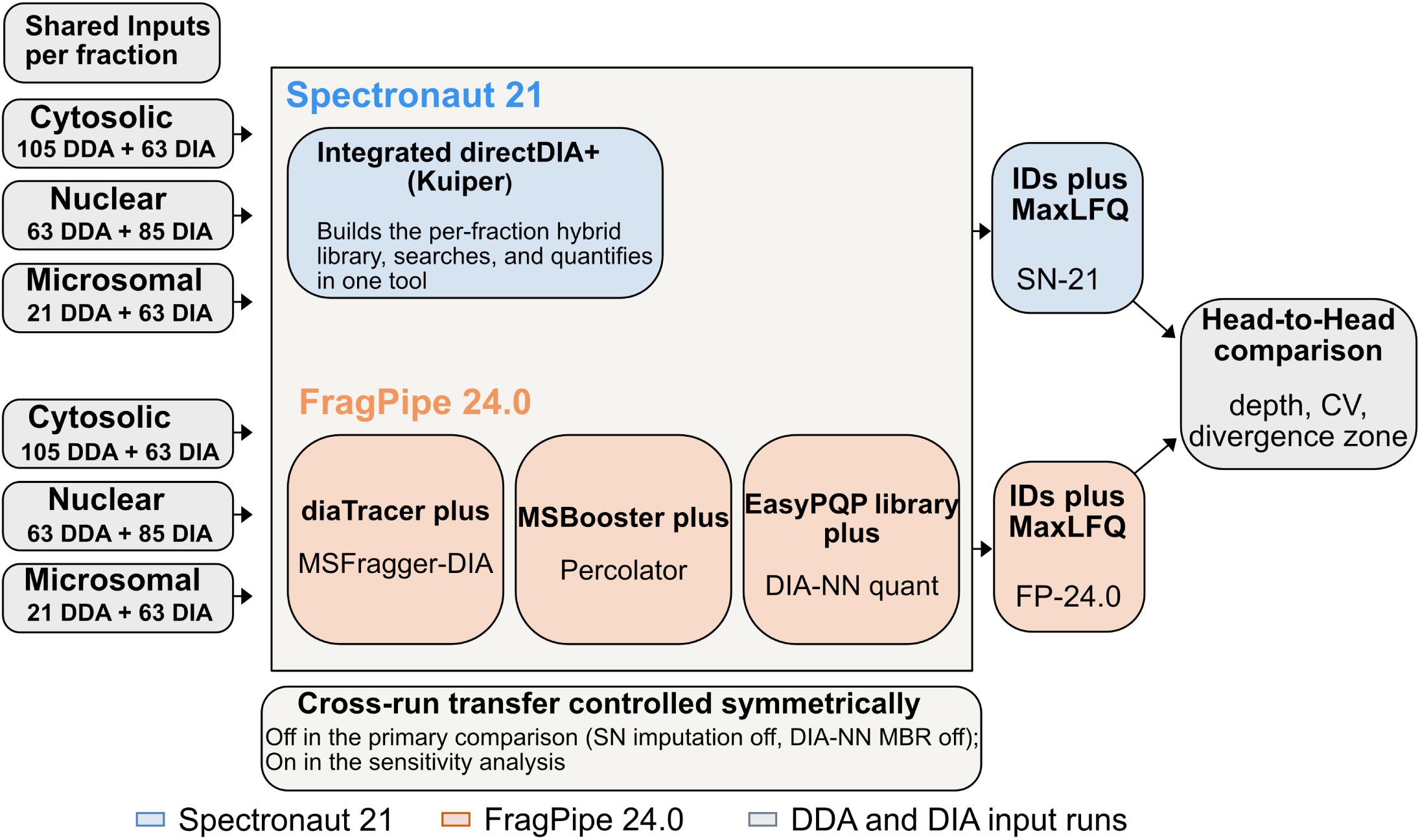
Confounder-controlled study design and per-fraction analysis workflow. Each subcellular fraction (cytosolic, nuclear, microsomal) was analyzed by both platforms from identical inputs, so that the only intended difference between the two workflows is the search-and-quantification engine. Shared per-fraction inputs (gray) comprise that fraction’s DIA runs and matched DDA runs (cytosolic, 105 DDA and 63 DIA; nuclear, 63 DDA and 85 DIA; microsomal, 21 DDA and 63 DIA) and feed both tracks. Spectronaut 21 (blue, upper track) runs an integrated directDIA+ (Kuiper) workflow that builds a per-fraction hybrid library and quantifies against it. FragPipe 24.0 (orange, lower track) runs a modular pipeline, with diaTracer pseudo-spectrum extraction and MSFragger-DIA search, MSBooster and Percolator rescoring, EasyPQP hybrid-library assembly, and DIA-NN quantification. Both platforms built their libraries the same way within each fraction and ran at matched parameters (1% FDR at protein and precursor level, strict trypsin cleavage, variable methionine oxidation, 20 ppm precursor and fragment tolerances, MaxLFQ roll-up). Cross-run transfer was controlled symmetrically, disabled on both platforms in the primary comparison (background-signal imputation off in SN-21, Match-Between-Runs off in DIA-NN) and enabled on both in the sensitivity analysis. Protein-group counts per platform per fraction are reported in Figure 2 and the Results.

#### Protein identifications and the divergence zone

We first compared the two engines at the protein-identification level. Because MSFragger-DIA and DIA-NN assign group-shared peptides differently from Pulsar and Kuiper, we expanded each group to its constituent UniProt accessions and compared the engines on accessions reported at a 1% protein-group FDR (Experimental Section). SN-21 identified more accessions than FP-24.0 in every fraction, and the margin widened from the cytosolic to the microsomal compartment, 7,106 versus 5,748 in cytosolic (+23.6%), 7,671 versus 6,000 in nuclear (+27.9%), and 7,156 versus 5,205 in microsomal (+37.5%) (Figure 2A). Much of each proteome was nonetheless shared, with 70 to 77% of each union recovered by both engines (Figure 2B; Supplementary Table S3). The remaining divergence zone was strongly asymmetric, SN-only accessions outnumbering FP-only by 10.6-to 18.3-fold, again largest in the microsomal fraction (Figure 2B,C; Supplementary Table S3). Because we matched library construction across fractions, this asymmetry reflects the search engines rather than library design.

**Figure 2.**
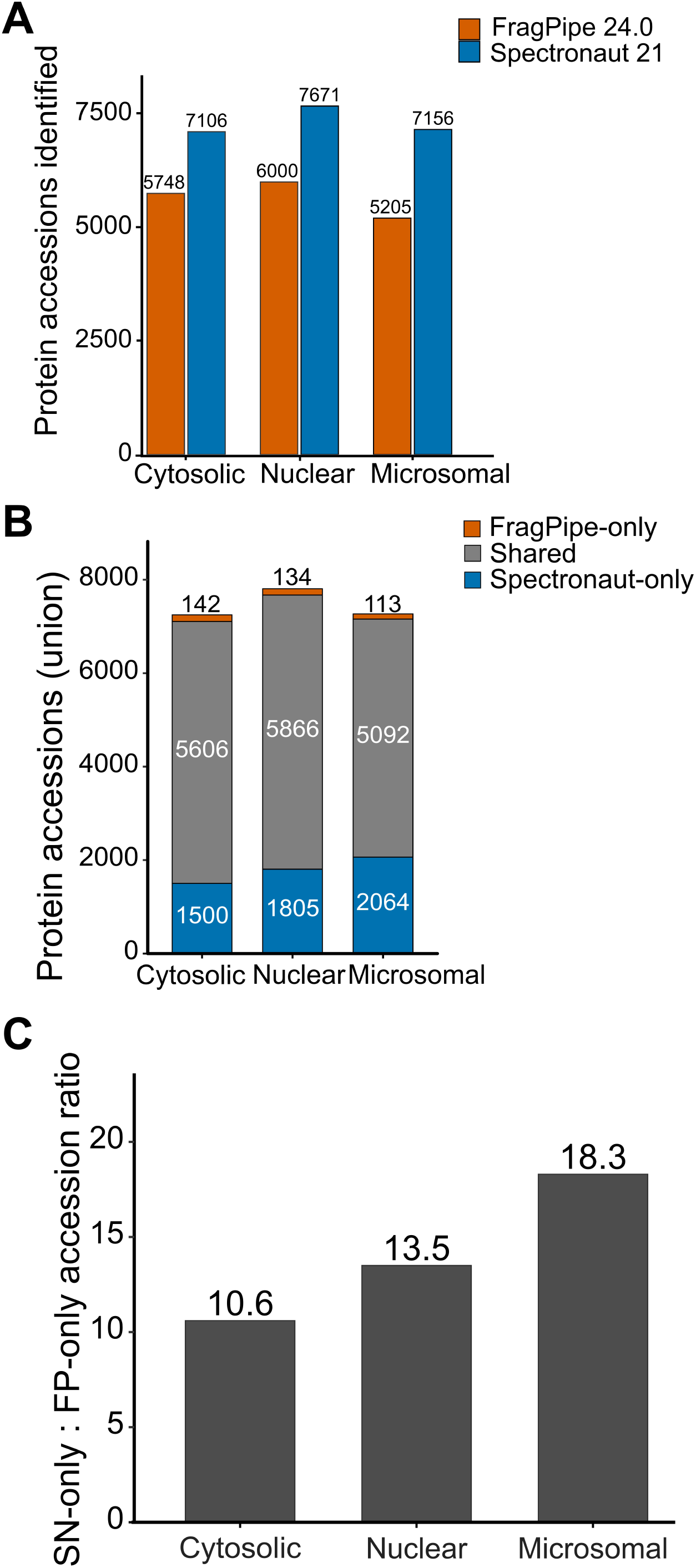
Headline identification metrics per fraction. (A) Protein accessions identified at 1% protein-group FDR by each platform in each subcellular fraction. SN-21 exceeds FP-24.0 in every fraction, by 23.6% in cytosolic, 27.9% in nuclear, and 37.5% in microsomal. (B) Composition of the union of both platforms per fraction, partitioned into accessions detected by both, by SN-21 only, and by FP-24.0 only. Accessions detected by both dominate every fraction at 70 to 77% of the union. (C) Ratio of SN-only to FP-only accessions per fraction. Because all three fractions were analyzed identically with fraction-restricted libraries, the ratio is not confounded by library construction; it is 10.6 in cytosolic, 13.5 in nuclear, and 18.3 in microsomal, largest in the microsomal fraction. Counts underlying all three panels are given in Supplementary Table S3.

#### Depth advantage under matched library construction

The SN-21 identification advantage is notable precisely because it persists after controlling for library construction. The advantage was largest in the microsomal fraction (Figure 3). Because each engine built its library the same way within every fraction, we cannot attribute the residual 23.6 to 37.5% advantage to library design. The findings reflect a property of the search engines themselves, and enabling cross-run transfer on both platforms left the ordering unchanged, with SN-21 ahead by 20.2%, 25.7%, and 35.8% across the three fractions (Supplementary Table S3).

**Figure 3.**
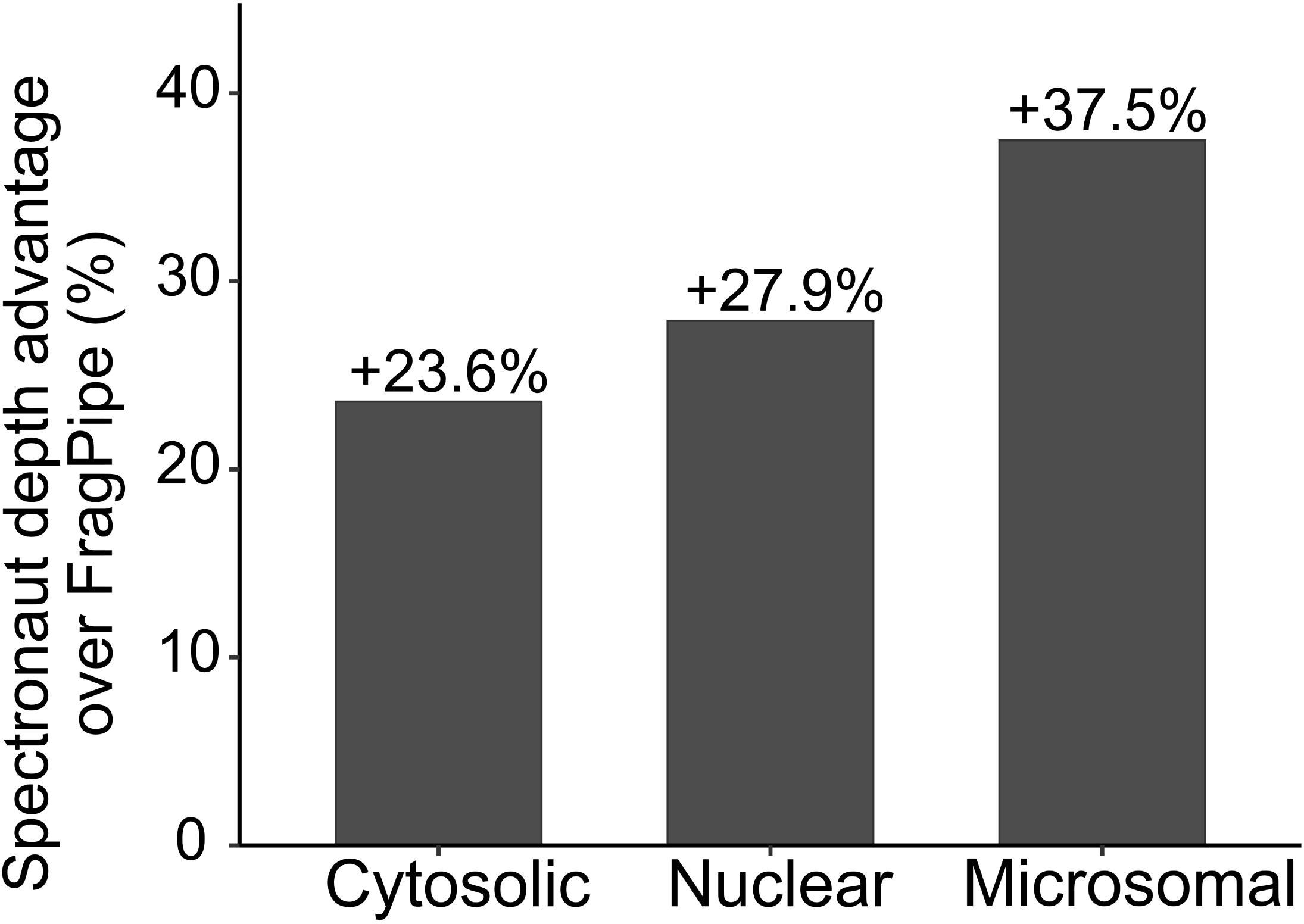
The depth advantage persists under matched library construction. SN-21 depth advantage over FP-24.0 in each subcellular fraction, expressed as the percentage by which SN-21 exceeds FP-24.0 in protein groups identified at 1% FDR. All three fractions were analyzed identically, with each platform building a fraction-restricted library from that fraction’s DDA and DIA runs, so differences in library construction do not confound the advantage. The advantage is substantial and consistent across fractions, 23.6% in cytosolic, 27.9% in nuclear, and 37.5% in microsomal, and largest in the microsomal fraction. We disabled cross-run transfer on both platforms. Figure 2A shows the underlying per-platform counts.

#### Concordant Quantification, but FP-24.0 Is the More Precise Quantifier Biological-replicate precision

We then shifted from what each engine identifies to how reproducibly it quantifies. We computed biological-replicate coefficients of variation (CVs) per protein group only for shared groups, so the comparison reflects quantitative behavior on jointly detected proteins rather than the divergence zone (Supplementary Figure S2). Here, the ordering reversed. FP-24.0 delivered tighter median CVs than SN-21 in every fraction (21.2 to 30.9% vs 25.9 to 37.3%), a gap of 4.1 to 6.4 percentage points that was consistent across compartments and robust to the cross-run normalization we enabled on both platforms (Figure 4A). This precision advantage is not an artifact of abundance coverage. When we stratified shared proteins by abundance, FP-24.0 was more precise than SN-21 in every abundance quartile of every fraction; rather than narrowing toward high abundance, the gap persisted or widened, reaching 4.1 to 6.3 percentage points among the most abundant proteins (Supplementary Table S11; Supplementary Figure S3). The two engines nonetheless agreed closely on the values they assigned to shared proteins (global Spearman 0.79 to 0.84; Figure 5), so FP-24.0’s advantage reflects reproducibility rather than a systematic difference in the signal each recovers.

**Figure 4.**
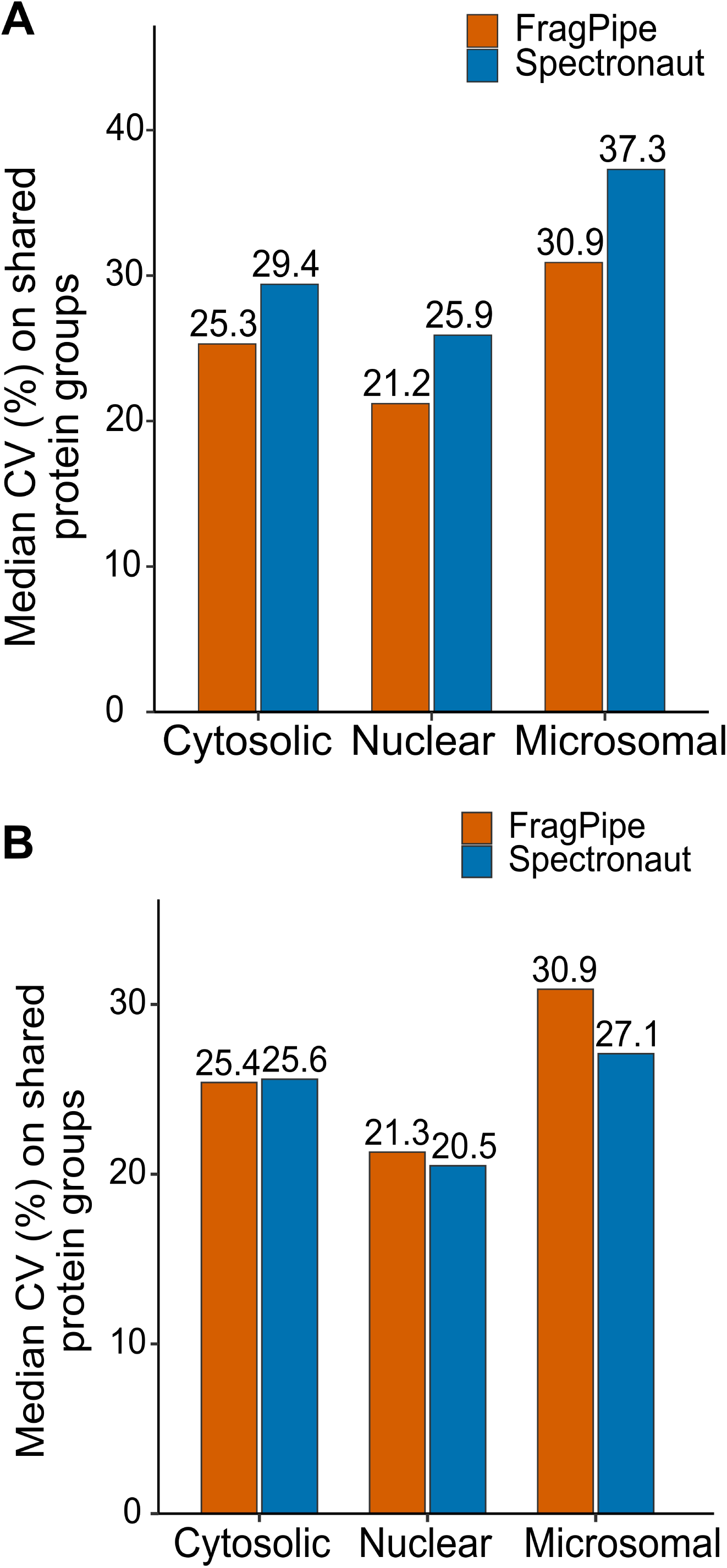
Coefficients of variation on shared protein groups. (A) Median coefficient of variation (CV) computed per protein group on shared protein groups only, per platform per subcellular fraction, with cross-run transfer disabled on both platforms. FP-24.0 delivers 4.1 to 6.4 percentage-point tighter CVs than SN-21 in every fraction. (B) The same comparison with cross-run transfer enabled on both platforms. SN-21’s background-signal imputation fills missing values with low-variance estimates and narrows or reverses the CV gap, an imputation effect rather than a genuine gain in per-precursor precision. We enabled cross-run normalization on both platforms throughout.

**Figure 5.**
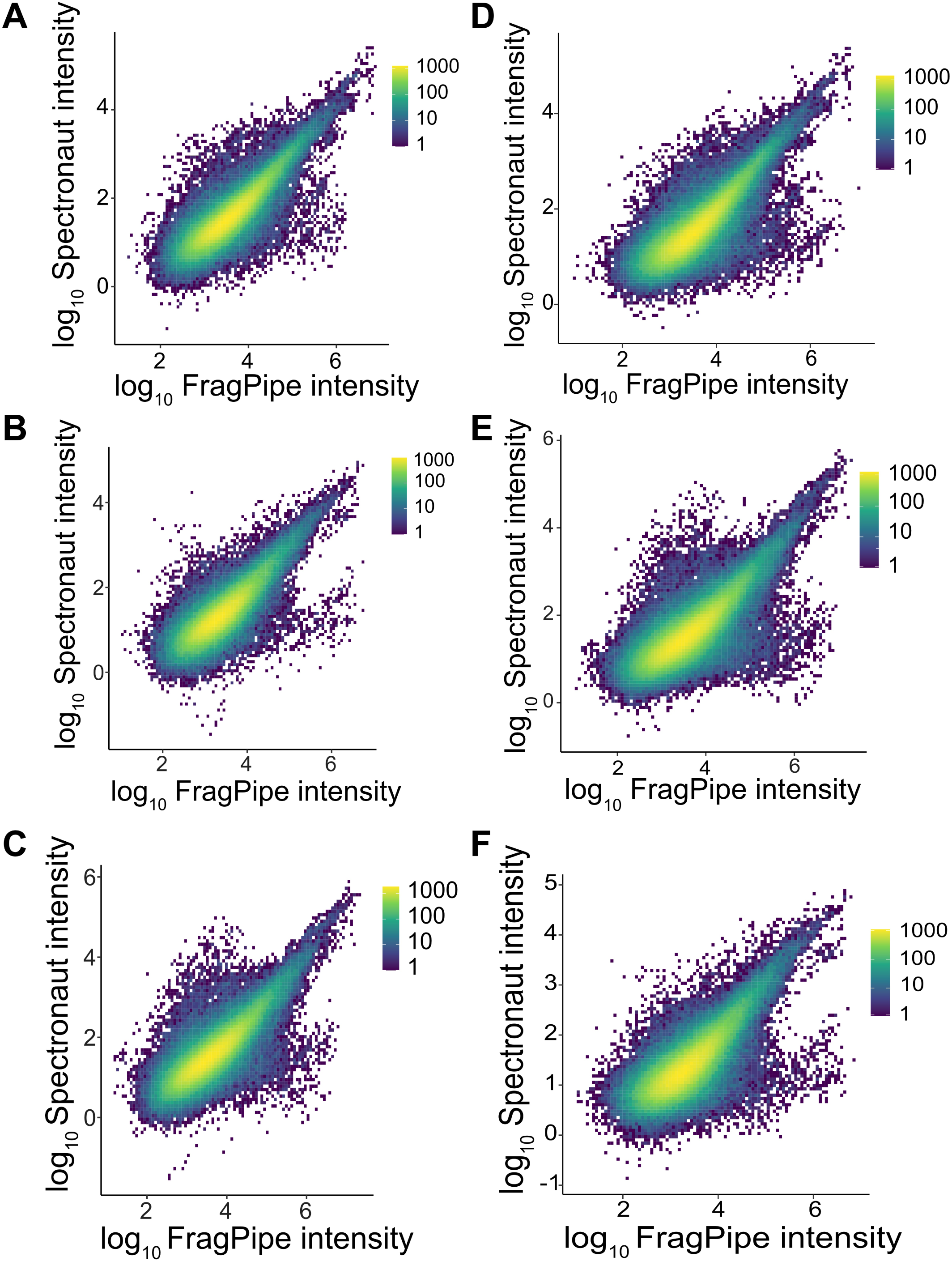
Cross-platform quantitative agreement on shared proteins across three fractions. Two-dimensional density of shared proteins on log_10_ MaxLFQ intensity, with FP-24.0 on the x axis and SN-21 on the y axis, where color gives the number of protein-by-run points per bin. (A to C) Cross-run transfer disabled on both platforms, showing the cytosolic, nuclear, and microsomal fractions from left to right. (D to F) Cross-run transfer enabled on both platforms (Match-Between-Runs in DIA-NN, background-signal imputation in SN-21), in the same three fractions from left to right. Because FP-24.0 and SN-21 report MaxLFQ on different absolute scales, offset by about 2 log_10_ units, the density ridge runs parallel to but below the one-to-one diagonal. With cross-run transfer disabled, the scale-invariant global Spearman rank correlations are 0.84 (cytosolic), 0.82 (nuclear), and 0.79 (microsomal), showing that the platforms rank shared-protein intensities almost identically. With cross-run transfer enabled, the correlations fall to 0.80, 0.78, and 0.74 respectively, so SN-21’s background-signal imputation does not improve cross-platform agreement. The composition of the shared and platform-specific accession sets is shown in Figure 2B. The divergence between platforms is concentrated in identification rather than in quantitation of jointly detected proteins.

**Figure 6.**
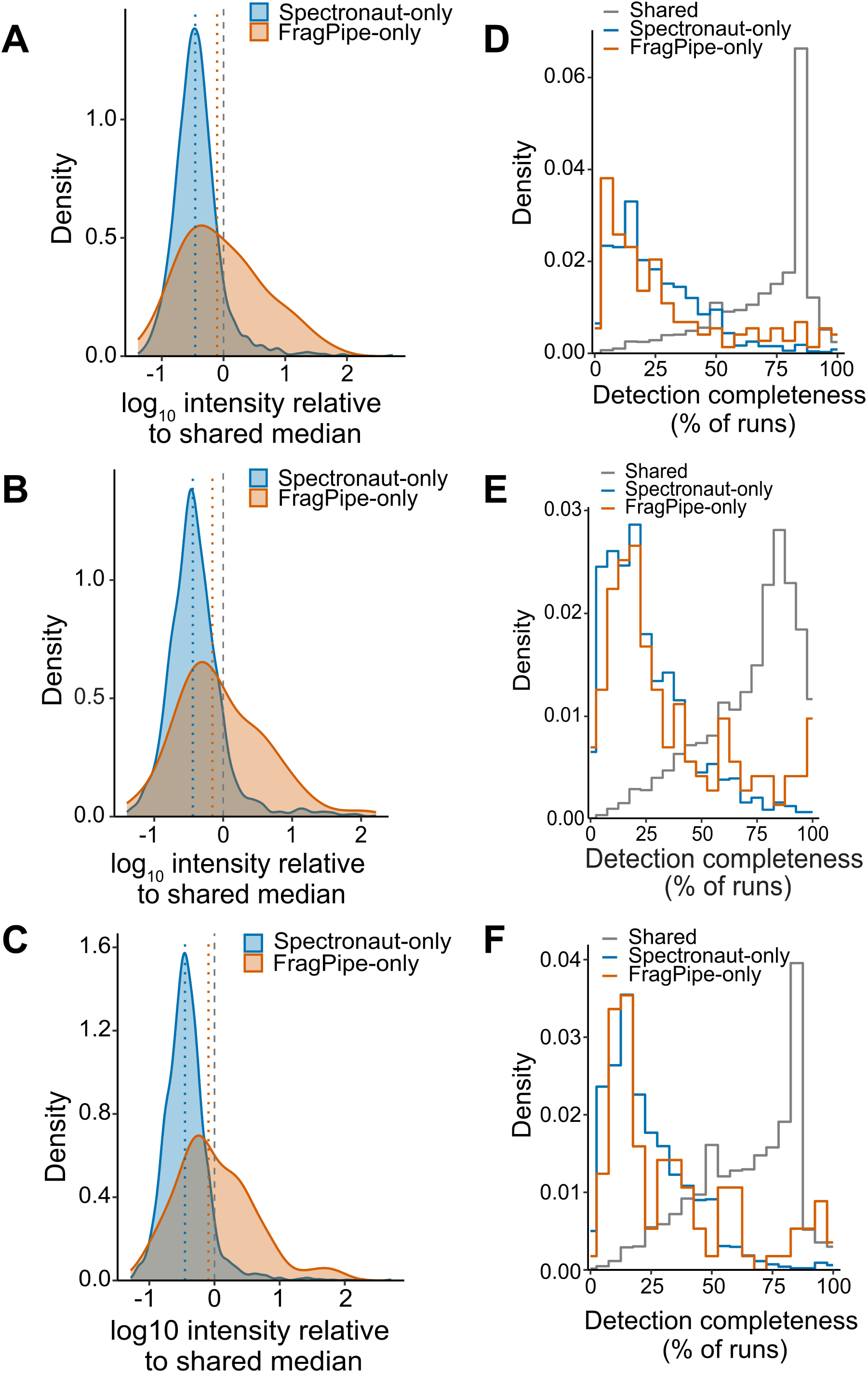
Mechanistic characterization of divergence-zone proteins. Row 1 (A to C) shows the distribution of median log_10_ protein intensity relative to the shared-proteome median, for SN-only (blue) and FP-only (orange) proteins in the cytosolic, nuclear, and microsomal fractions. Each protein is expressed relative to its own platform’s shared median, so both divergence zones share one axis despite the different absolute FP-24.0 and SN-21 intensity scales, and the gray dashed line at zero marks the shared baseline. Both zones peak below the shared median, and the SN-only zone sits deeper than the FP-only zone, with median shifts of 0.46 and 0.10 (cytosolic), 0.44 and 0.16 (nuclear), and 0.45 and 0.09 (microsomal) log_10_ units below the shared median. The shallower FP-24.0 shift is consistent with FP-24.0 being the more conservative platform. Dotted vertical lines mark each zone’s median. Row 2 (D to F) shows per-run detection completeness (percent of runs where a protein was quantified) for shared (gray), SN-only (blue), and FP-only (orange) proteins. Both divergence zones are dominated by proteins detected in fewer than half of the runs, whereas shared proteins show a strong secondary peak at high completeness. Distributions are drawn as step outlines so no two overlap into an unkeyed color. Together the low intensity and the intermittent detection are consistent with the divergence-zone proteins being low-abundance detections at the sensitivity edge of DIA-MS rather than noise-driven false positives.

#### Shared-proteome agreement and cross-run transfer

Focusing on the proteins both engines detected (5,092 to 5,866 per fraction; Figure 2B), we found strong, scale-independent quantitative agreement. Global cross-platform Spearman correlations on shared MaxLFQ intensities were 0.84, 0.82, and 0.79 in the cytosolic, nuclear, and microsomal fractions, respectively, and held across the intensity range (Figure 5A-C). Enabling cross-run transfer eroded this concordance, the cross-platform Spearman correlation falling to 0.80, 0.78, and 0.74 (Figure 5D-F). Had transfer genuinely sharpened SN-21’s measurements, its values would agree with FP-24.0’s independent measurements of the same proteins at least as well as before. Instead the agreement fell, which shows that SN-21’s background-signal imputation lowers the apparent replicate CV (Figure 4B) by filling missing values with low-variance estimates that diverge from FP-24.0’s measured values. The tighter CV under transfer therefore reflects imputation rather than a real gain in precision. For proteins both engines detect, they assign closely matching quantities, so the two platforms differ in which proteins they detect, not in the values they assign to the proteins they share.

### The Divergence Zone Is a Reproducible Low-Abundance Boundary with Distinct Biology

#### Intensity and detection completeness

We next asked whether proteins detected by only one platform, its divergence zone, are genuine low-abundance proteins at the edge of detection or simply noisy, marginal identifications. For each platform’s divergence zone, we characterized both the intensity of those proteins and their detection completeness, the number of runs in which each protein was quantified, relative to that platform’s shared-proteome baseline (Figure 6).

Both measures placed the divergence zone at the sensitivity edge rather than in the noise. SN-only proteins sat at the low-intensity margin of SN-21’s range, roughly 2.8-fold fainter than shared proteins and within the bottom 16 to 19% of the intensity distribution (Figure 6). They were also detected intermittently, quantified in a median of 13 to 18 of the 63 to 85 runs per fraction, compared with 43 to 66 for shared proteins, and only 7.7 to 12% appeared in more than half the runs, compared with 73 to 83% of shared proteins. FP-only proteins showed the same faint, intermittent pattern within FP-24.0 (Figure 6).

This pairing of faint signal and sporadic detection is the expected signature of genuine low-abundance proteins sampled near the Poisson-limited detection threshold, rather than noise-driven false positives, which would instead surface as random single-run hits (Figure 6). Two further observations support this interpretation. First, the FP-only zone fell below its shared baseline by a shallower margin than SN-only (0.09 to 0.16 log_10_ units across the three fractions), consistent with FP-24.0 reaching less deeply into the low-abundance regime rather than detecting a qualitatively different population (Figure 6). Second, neither divergence zone was enriched for common contaminants (Supplementary Figure S1). The SN-only contaminant fractions (0.83 to 1.20%) tracked the shared baseline (0.69 to 0.98%), and although the much smaller FP-only sets ran higher (1.49 to 4.23%, driven by keratins and immunoglobulins), contaminants remained a small minority throughout. Together, the faint-but-real intensity profile and the low contaminant content indicate that each platform’s divergence zone is dominated by genuine low-abundance proteins at the sensitivity edge of DIA rather than by contaminants or noise.

#### Subcellular-compartment composition

We next asked whether this divergence reshaped the proteome as a whole or stayed confined to its low-abundance margin. Mapped to gene symbols, both engines recovered most of each fraction’s proteome, with roughly three-quarters shared (70 to 77%; Supplementary Table S3). The remainder was modest and lopsided, with SN-21 recovering genes that FP-24.0 missed an order of magnitude more often, and vice versa.

The two proteomes also matched in subcellular-compartment composition. Scoring each full proteome by the percentage of its genes annotated to each subcellular compartment, the SN-21 and FP-24.0 proteomes were nearly superimposable, agreeing to within about five percentage points in every compartment and fraction (Figure 7). The extra proteins SN-21 detects are therefore distributed across compartments in the same proportions, so its greater depth does not come from over-sampling any one compartment. The functional divergence between the platforms is therefore confined to their divergence-zone sets rather than spread across the proteome as a whole.

**Figure 7.**
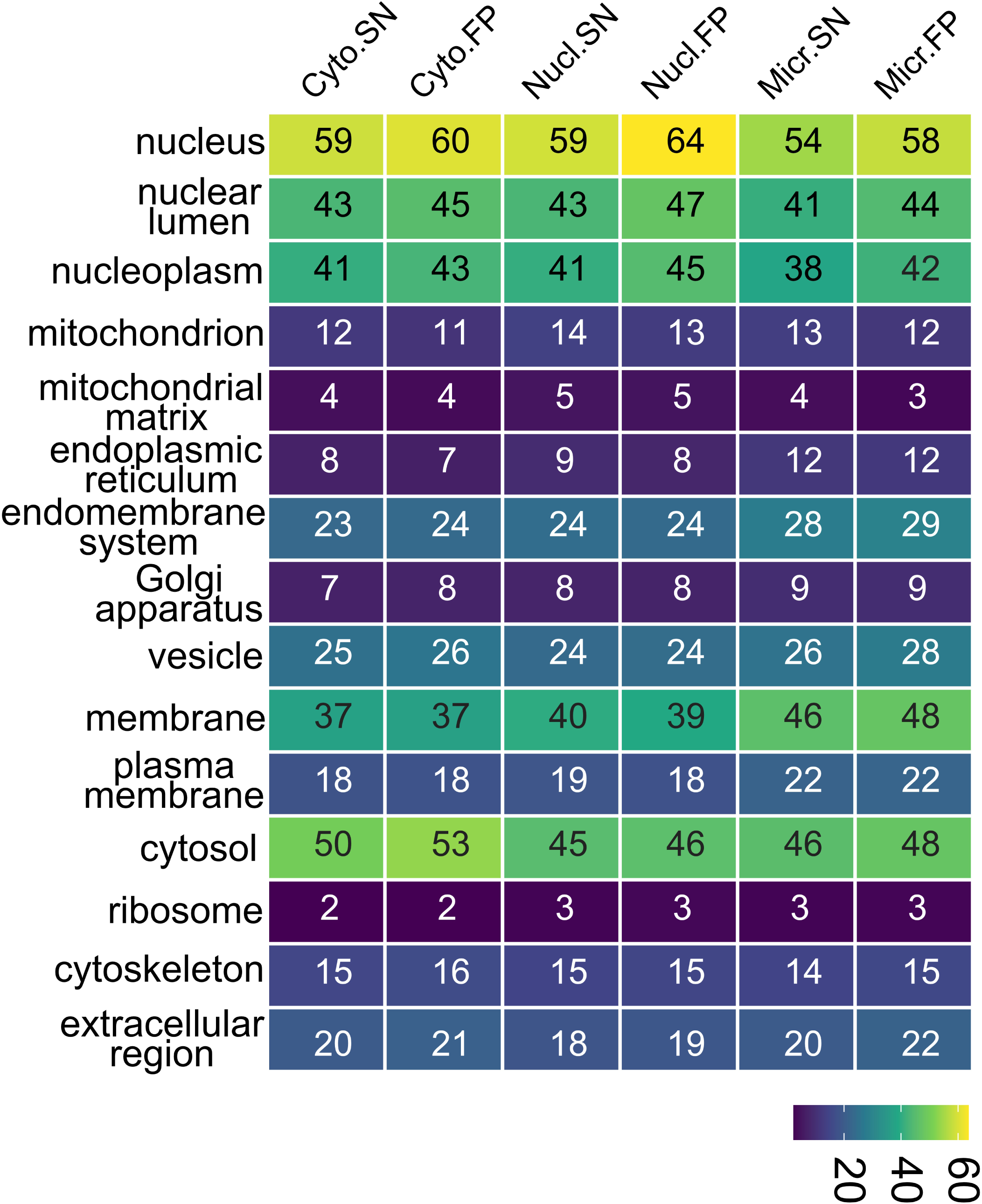
Whole-proteome compartment composition of the two platforms. Percentage of each platform’s full detected proteome annotated to each subcellular compartment (Gene Ontology cellular component), for SN-21 and FP-24.0 in the cytosolic, nuclear, and microsomal fractions, computed as membership against the annotated human genome. The two compositions are nearly identical in every compartment and fraction, differing by at most a few percentage points, showing that the platforms recover functionally equivalent whole proteomes and that their functional divergence is confined to the platform-specific divergence zone.

#### Divergence-zone functional enrichment

To test whether the divergence zone is biologically structured or a random shortfall, we ran functional enrichment on the SN-only and FP-only gene sets in each fraction, using that fraction’s measured proteome as background (Figure 8, Supplementary Table S4). Enrichment separated the two divergence zones into distinct biology rather than one shared signal sampled at different rates. The zones are also very unequal in size, with FP-24.0 contributing only 140, 133, and 113 unique genes across the three fractions versus 1,492, 1,798, and 2,057 for SN-21, so SN-21’s greater depth extends into the divergence zone as well. The platforms differ by the identity of the enriched terms rather than their number.

**Figure 8.**
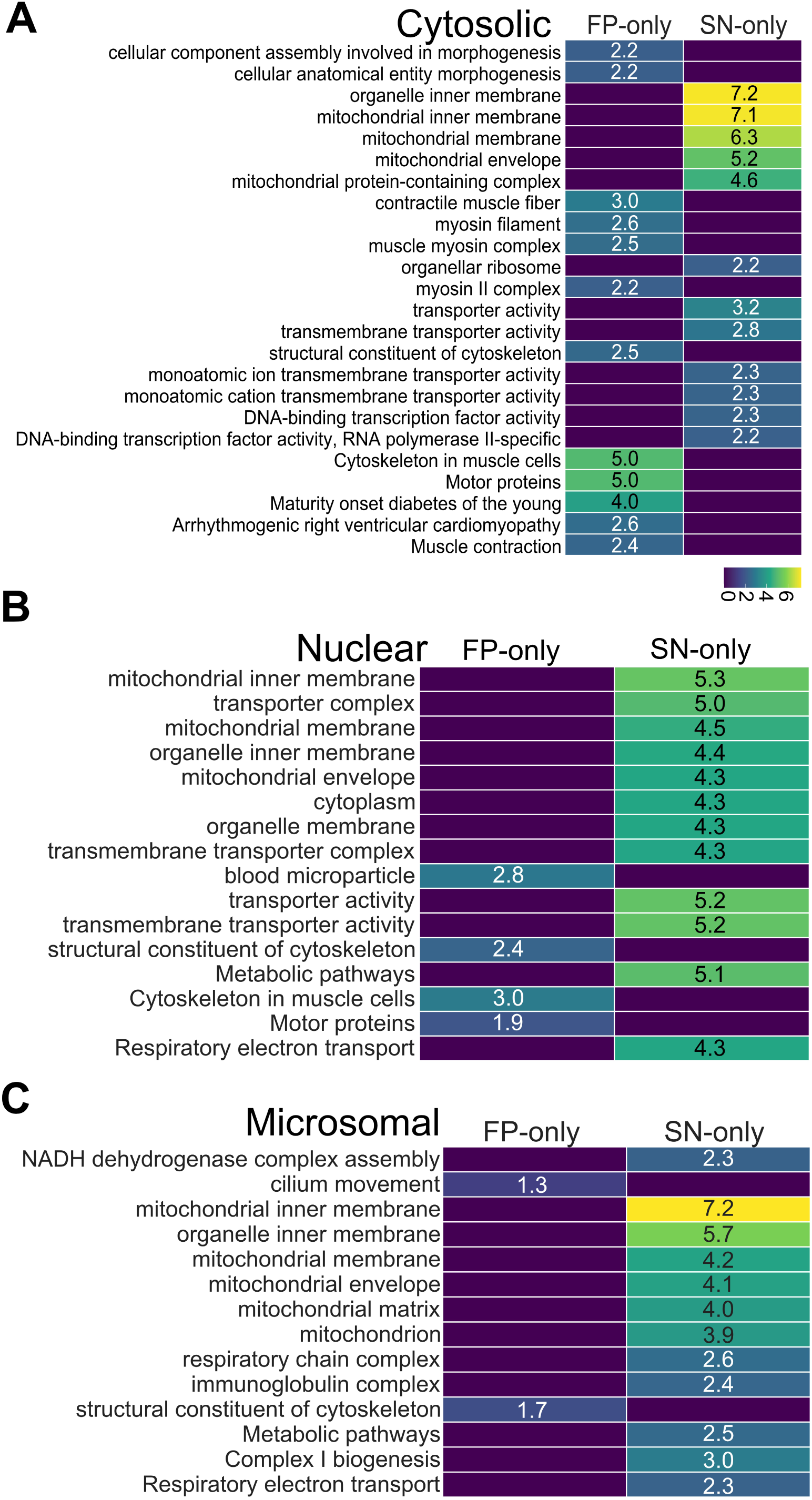
Divergence-zone GO term enrichment across fractions and platforms. Three separate per-fraction panels (A cytosolic, B nuclear, C microsomal), each a two-column heatmap comparing the FP-only divergence zone (left column) and the SN-only divergence zone (right column) in that fraction. Rows are GO, KEGG, and Reactome terms enriched (Benjamini-Hochberg FDR < 0.05, no IEA) in either divergence zone, showing up to the twelve most significant terms per column and sorted by term source and then by best p-value. Cell color is the -log_10_ FDR from g:Profiler on a viridis scale shared across all three panels, so magnitudes are directly comparable, and each cell also carries its numeric -log_10_ FDR. A value of zero, shown at the dark bottom of the scale, means the term is not significant in that column. The full significant-term counts are 47 and 33 (cytosolic), 4 and 82 (nuclear), and 2 and 29 (microsomal) for the FP-only and SN-only sets respectively, with the complete tables in Supplementary Table S4. The key observations are (i) the strongest enrichments in the study are the Spectronaut-only mitochondrial terms, which reach -log_10_ FDR near 7 in the cytosolic and microsomal fractions; (ii) mitochondrial biology recurs across all three SN-only columns, most prominently in the cytosolic and microsomal fractions, alongside membrane-transport terms and, in the cytosolic fraction, DNA-binding transcription-factor activity; (iii) the FP-only columns are dominated by cytoskeletal, motor-protein, and muscle terms in the cytosolic and nuclear fractions; and (iv) the microsomal FP-only column is nearly empty, reaching significance for only two weak cytoskeletal terms, so the microsomal divergence zone is overwhelmingly Spectronaut-only.

Across all three fractions, the SN-only set was dominated by mitochondrial biology. In the cytosolic fraction, mitochondrial inner-membrane, membrane, and envelope terms led the enrichment, and in the microsomal fraction the strongest SN-only terms were again mitochondrial, spanning the inner membrane, the matrix, and respiratory-chain and Complex I biogenesis (Figure 8, Supplementary Table S4). The cytosolic set carried one further signal of note, a significant enrichment for DNA-binding transcription factors and their RNA polymerase II subclass, specific to the cytosolic fraction and absent from the nuclear and microsomal SN-only sets. Because the AR bait is itself a transcription factor, that cytosolic enrichment is biologically pertinent, even though the dominant SN-only theme across every fraction is mitochondrial.

In all three fractions, the FP-only set was instead enriched for cytoskeletal and motor-protein biology. The signal was strongest in the cytosolic fraction, which generated the most terms (47), spanning muscle cytoskeleton, motor proteins, contractile fibers, and cytoskeletal structural components, with muscle-disease terms drawing on the same proteins (Figure 8, Supplementary Table S4). The nuclear FP-only set carried the same cytoskeletal and motor signature on fewer terms, and the microsomal FP-only set, the smallest of the three, reached significance for only two weak terms, reflecting an FP-only set too small to contribute functional signal of its own, so the microsomal divergence zone is overwhelmingly an SN-21 phenomenon.

The cytoskeletal FP-only genes warranted a closer look, because they are dominated by tissue-restricted family members not expected in an epithelial culture, ACTA1 (skeletal muscle alpha-actin), ACTC1 (cardiac alpha-actin), the myosin heavy chains MYH11 and MYH13, the tissue-restricted tubulins TUBA8 and TUBB1, the corneal keratin KRT12, and the keratin pseudogene KRT87P. Because these paralogues share most of their tryptic peptides, their identifications could in principle rest on shared peptides misassigned within a family rather than on genuine detection. We addressed this at two levels. At the gene level, we define platform-specific sets throughout, so a gene the two engines report under different accessions counts as shared, ruling out any artifact of accession bookkeeping. At the peptide level, we tested the FP-only identifications directly, asking whether each gene in the FP-24.0 precursor output carries a proteotypic peptide that maps uniquely to it in the search database (Supplementary Table S10). Seven of the eight, ACTA1, MYH11, MYH13, TUBA8, TUBB1, KRT12, and KRT87P, carried quantified proteotypic peptides, so their FP-only identifications rest on unique evidence rather than homology inference. The lone exception was ACTC1, supported only by peptides shared with the expressed actins. The cytoskeletal FP-only signal therefore reflects genuine recovery of low-abundance, tissue-restricted proteins that SN-21’s more conservative inference does not report, with ACTC1 the single inference artifact.

The complete separation between the two divergence zones is more revealing than any single enriched term. Across all three fractions SN-21’s divergence zone was enriched for mitochondrial biology and membrane transport, and in the cytosolic fraction for DNA-binding transcription factors, whereas FP-24.0’s was enriched for cytoskeletal, motor-protein, and muscle biology. The two zones never shared a functional class, so the engines were surfacing systematically different protein classes rather than sampling the same biology at different rates. The most economical explanation is that divergence-zone proteins are low-abundance detections at the sensitivity edge of DIA, populated by different functional classes on each engine, rather than platform-agnostic noise drawn by different algorithms.

#### Cross-fraction recurrence

A recurring theme need not mean recurring proteins, since different proteins could produce the same functional class in each fraction, making the bias a property of the sample rather than the engine. We therefore asked whether the same individual genes, not merely the same themes, fall into a given engine’s divergence zone in all three fractions. They do, and asymmetrically, with 146 genes recurring in the SN-only set across every fraction against 10 in the FP-only set, mirroring the per-fraction SN-only-to-FP-only imbalance (Supplementary Table S7). Each engine thus has its own reproducible blind spot, the 146 genes only SN-21 recovers and the 10 only FP-24.0 recovers, and these same genes fall outside the other engine in every fraction. These recurring genes are also the same protein classes as the full divergence zones, mitochondrial and membrane transport for SN-21, and cytoskeletal and motor for FP-24.0 (Supplementary Table S5). The bias is a fixed property of the analysis platform, not an artifact of any one fraction’s composition.

### SN-21 Recovers More of the Known AR Interactome, at Its Low-Abundance Margins

#### Recovery of the known AR interactome

In discovery proteomics the search engine gatekeeps what can be found, since its reach over both known and novel members of a network sets how far a study can push past established biology. A platform’s blind spot therefore matters only where it falls on the biology that study is trying to map. Our dataset answers this directly, because its target, AR, is defined independently of any search engine by a curated 989-member interactome. Intersecting the per-fraction shared and divergence-zone sets with that reference (Supplementary Table S6), both engines recovered the great majority of it, 852 of 989 members in the cytosolic fraction (86.1%), 888 in the nuclear (89.8%), and 851 in the microsomal (86.0%), the bulk of them, 798, 827, and 759 respectively, detected by both (Figure 9A). The known interactome therefore lives predominantly in the shared proteome, and neither engine defines it alone.

**Figure 9.**
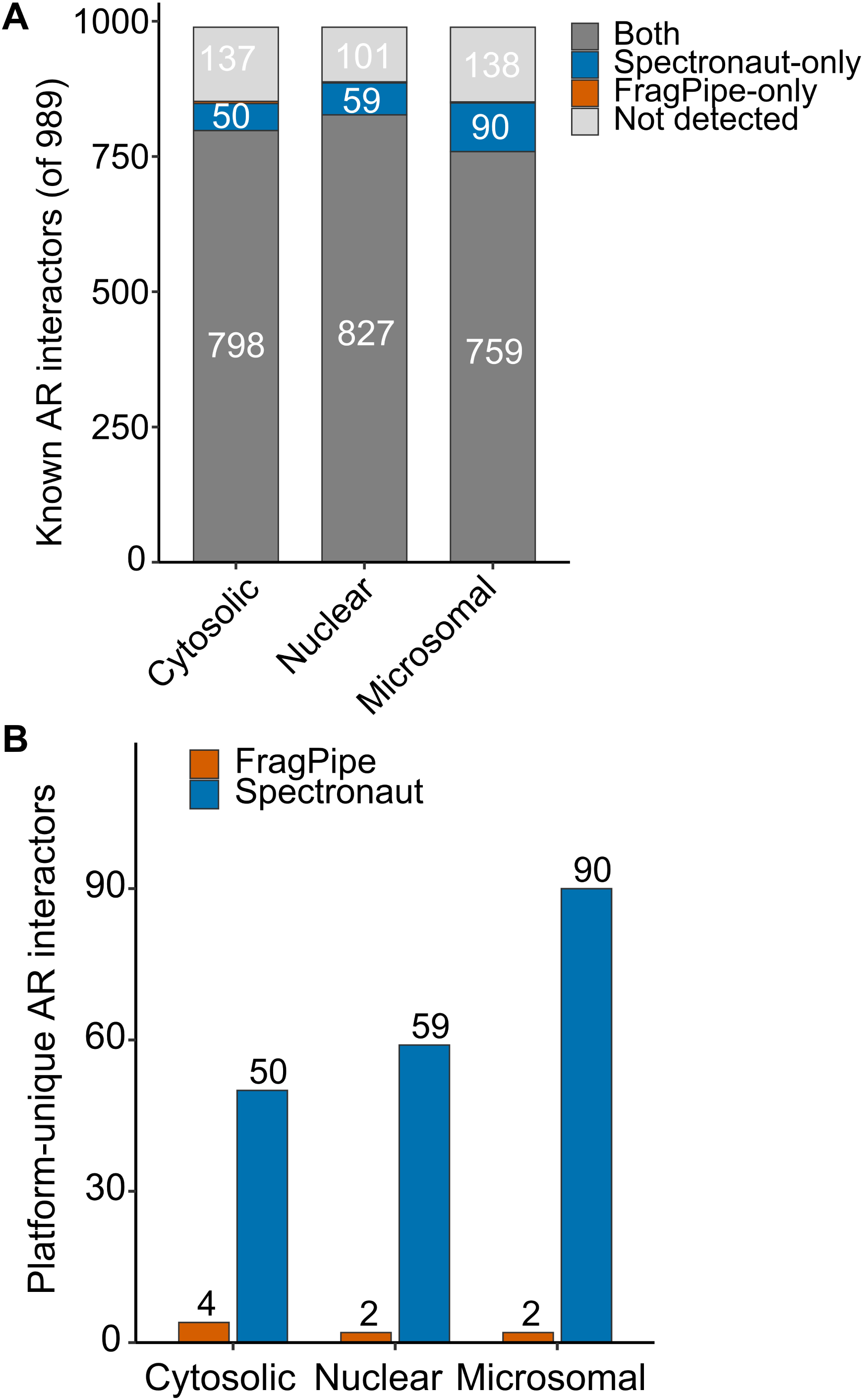
Recovery of the known AR-interactome by each search method. (A) Fate of the 989 known AR-interactome members in each subcellular fraction, shown as stacked bars of members detected by both platforms (gray), detected only by SN-21 (blue), detected only by FP-24.0 (orange), and not detected in that fraction (light gray). Between 86.0% and 89.8% of the interactome is recovered per fraction and the large majority is common to both platforms. (B) Members recovered by only one platform, per fraction. SN-21 uniquely recovers 50, 59, and 90 members in the cytosolic, nuclear, and microsomal fractions, against 4, 2, and 2 for FP-24.0.

With most of the known interactome shared between the engines, the question that separates them is how much known AR biology each recovers in its divergence zone, the platform-specific part where they disagree. There the recovery was strongly asymmetric. Within a fraction, SN-only uniquely recovered 50, 59, and 90 interactome members across the cytosolic, nuclear, and microsomal fractions against 4, 2, and 2 for FP-only (Figure 9B, Supplementary Table S6). Counting only members that one engine detected and the other never did in any fraction, 34 were unique to SN-21 and 3 to FP-24.0 (GLI2, AURKB, and UBB). Independent PubMed co-citation pointed the same way, with the SN-21-unique members spanning 545 androgen receptor papers against 23 for the FP-24.0-unique members (per-member counts in Supplementary Table S6). In keeping with the low-abundance character of the divergence zone, the SN-21-unique members were largely functional co-regulators and pathway components rather than the abundant canonical interactors both engines detect, among them the aryl hydrocarbon receptor AHR, a documented physical AR partner. The most-cited FP-24.0-unique member, GLI2, carried 13 papers.

#### The platform-unique AR network

To understand which biological pathways each platform’s unique members represent, we built a physical-association network of the strictly platform-unique members in Cytoscape (Figure 10) and tested functional enrichment on the broader SN-only interactome set (Supplementary Table S9). The 152 SN-only members returned coherent enrichment led by KEGG Pathways in cancer (23 of 253 term proteins, FDR 1.4 × 10_−9_), whereas the FP-only members were too few to support an equivalent analysis.

**Figure 10.**
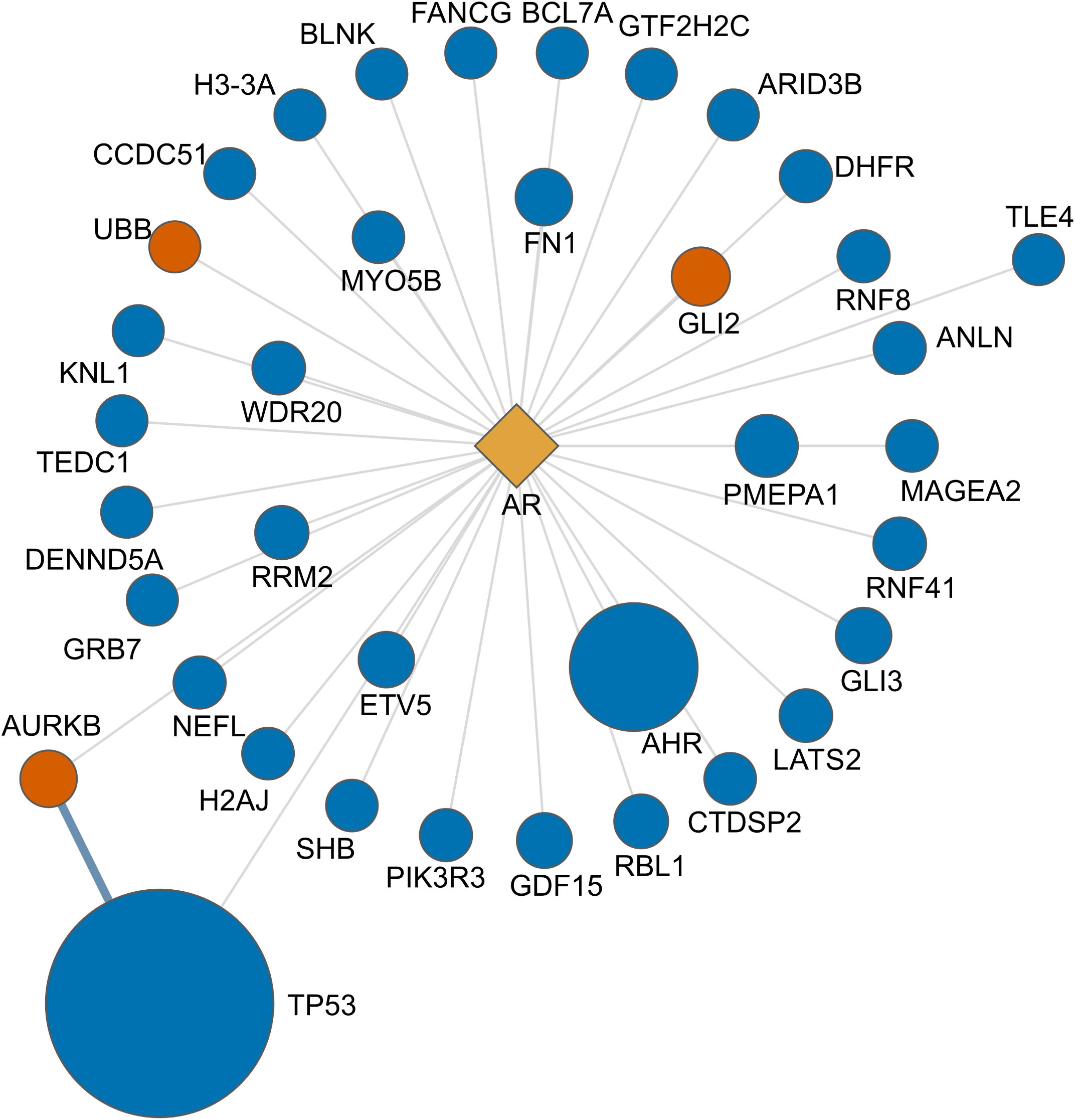
Physical-interaction network of the platform-unique AR-interactome members. The androgen receptor bait (gold) is shown with the 34 known AR-interactome members recovered only by SN-21 (blue) and the 3 recovered only by FP-24.0 (orange), where platform-unique is defined strictly as detection by one platform in at least one fraction and by the other in no fraction. Thin gray edges connect the bait to each member, which is an AR-interactor by definition, and steel edges, where present, are STRING physical interactions between members at a confidence score of 0.70 or above. Node area scales with the number of PubMed papers co-citing the member with the androgen receptor. Node lists, co-citation counts, and enrichment are provided in Supplementary Table S9.

SN-21 recovered more of the known AR interactome than FP-24.0 because it identified more proteins overall (Figure 3). The extra members sit at the low-abundance edge of the divergence zone (Figure 6), where only the deeper search reaches them. For a proximal-interactome study, whose aim is to enumerate a bait’s direct and indirect partners, the engine that recovers more of the known interactome recovers more of the biology, and on this AR dataset that engine is SN-21, which uniquely recovered 34 known members against 3 and carried far more of the associated literature (545 against 23 papers). FP-24.0 holds the complementary advantage in quantitative precision, with median CVs 4.1 to 6.4 percentage points tighter. On this dataset, then, SN-21 is the more sensitive engine for recovering the AR interactome and FP-24.0 the more precise quantifier.

## Discussion

### FP-24.0 Is the More Precise Quantifier, across the Dynamic Range

FP-24.0 quantified shared proteins more reproducibly than SN-21 in every fraction, by 4.1 to 6.4 percentage points of median CV. We first asked whether this gap was an artifact of dynamic-range coverage, since SN-21 reaches further into the low-abundance regime where counting statistics inflate replicate variation. Stratifying the shared proteins by abundance ruled that out. FP-24.0 was the more precise quantifier in every abundance quartile of every fraction, and rather than closing toward high abundance, the gap persisted or widened, reaching 4.1 to 6.3 percentage points among the most abundant, best-measured proteins (Supplementary Table S11, Supplementary Figure S3). The two engines nonetheless agreed closely on the values they assigned to shared proteins (Spearman 0.79 to 0.84), so the difference reflects reproducibility rather than the central estimate. FP-24.0 therefore holds a genuine, algorithm-level precision advantage rather than a coverage artifact. We do not pinpoint its origin here. The two quantifiers differ in how they select quantifying precursors, handle interference, and roll peptides up to proteins, and attributing the gap to a single step would require each vendor’s internals. Wherever downstream inference depends on replicate reproducibility, the advantage is real and measurable.

### Matched Configuration as a Methodological Contribution

A second contribution is methodological. The two platforms handle cross-run transfer differently, but neither enables it by default. The FragPipe-packaged DIA-NN leaves Match-Between-Runs off by default, and Spectronaut 21 has no MBR equivalent, providing background-signal imputation only as an optional quantification-stage step that is off unless enabled. Any depth advantage measured with transfer active conflates three things, the search engine and its library-generation stack, the peptide-level quantifier, and the cross-run rescue layer. We separated them by controlling library construction directly and setting cross-run transfer symmetrically, off on both platforms and then on. The depth ordering held at both settings, so SN-21’s advantage did not depend on asymmetric feature use. For head-to-head benchmarks, a matched configuration of this kind, controlled libraries and symmetric transfer, is a defensible reporting standard, because it removes the two systematic asymmetries that otherwise dominate depth comparisons. We recommend it for future benchmarking.

### Controlling Library Construction Reveals an Engine-Level Depth Difference

A third contribution is that controlling library construction exposes an engine-level depth difference. Each platform built a fraction-restricted library from that fraction’s own DDA and DIA runs, so library construction was identical for both within each fraction. Under this matched design the SN-21 depth advantage was substantial and consistent in all three fractions and largest in the microsomal (23.6%, 27.9%, and 37.5%). This reframes the platform-comparison literature. Depth claims built on vendor defaults, mixed libraries, or different analysis modes conflate library and transfer choices with the engine itself. When both are matched, as here, the engine-level difference between SN-21 and FP-24.0 is larger, not smaller, than confounded comparisons imply, a consistent property of the two platforms rather than an artifact of library design.

### Implications for AR-Interactome Biology

For laboratories using DIA to map the AR proximal interactome, the recovery asymmetry has direct consequences. Counting only interactome members that one platform detected while the other never did in any fraction, 34 were unique to SN-21 and 3 to FP-24.0 (Figure 10). The same asymmetry held within each fraction, with SN-21 uniquely recovering 50, 59, and 90 members versus 4, 2, and 2 (Figure 9). Because we matched library construction, this is not a library-construction advantage. The SN-unique members also carried far more androgen receptor literature, 545 co-citing papers against 23, and, consistent with the low-abundance character of the divergence zone, they were largely functional co-regulators and pathway proteins rather than the abundant canonical interactors both platforms detect. Several have direct experimental ties to AR in the primary literature. The aryl hydrocarbon receptor AHR is a documented physical AR partner (40) and the Hedgehog effector GLI3 physically interacts with AR (41), while PMEPA1 recruits the NEDD4 ligase to drive AR degradation (42) and WDR20 stabilizes AR as a subunit of the USP12-UAF1-WDR20 deubiquitinase (43). On the FP-unique side, the most-cited member, GLI2, likewise physically interacts with AR (44) and carried 13 co-citing papers. A study run on FP-24.0 alone would therefore omit an identifiable, non-random, literature-associated slice of the AR interaction landscape. Transcription-factor recovery is a concrete instance. In the cytosolic fraction, DNA-binding transcription factor activity was a significant SN-only enrichment (GO:0003700, 141 of 517 term proteins; Figure 8, Supplementary Table S4). Transcription factors are low-copy regulatory proteins at the low-abundance edge of the dynamic range (Figure 6), so they likely fall below FP-24.0’s more conservative detection threshold while remaining within SN-21’s deeper reach. For studies where the surrounding cell biology is the question, that is a real gap in what can be detected, not a marginal edge effect.

### A Generalizable Principle of Complementary Biology

The magnitudes here are particular to this AR dataset, but the cross-fraction concordance (Figure 9) shows platform bias recurs across all three fractions rather than being confined to one, pointing to a more general principle. One hundred and forty-six genes recur in the SN-only divergence zone across all three fractions, and 10 in the FP-only zone. The recurrent themes match the aggregate ones, mitochondrial biology and membrane transport on the SN-21 side, with DNA-binding transcription factors a further cytosolic signal, and cytoskeletal and muscle biology on the FP-24.0 side (Figure 8, Supplementary Tables S5 and S7). The bias is therefore a property of the platforms, not a fraction-specific quirk, and it is localized rather than diffuse. At the whole-proteome level the two are compositionally equivalent, agreeing to within about five percentage points in every compartment (Figure 7), so the divergence lives in the platform-specific sets, not the bulk proteome. For discovery-driven work aiming to enumerate a full set of putative interactors or a full functional profile at high sensitivity, running both platforms is defensible. They are better understood as complementary probes with systematically different sensitivity profiles than as redundant tools expected to agree. What one platform misses, the other catches, and the missed biology is systematic, literature-validated, and biologically meaningful rather than random or cosmetic.

### Limitations

Several limitations should be acknowledged. First, this study is anchored on a single dataset acquired on a single instrument (Bruker timsTOF Pro2) from a single cell system (LNCaP stably expressing APEX2-AR), using a single acquisition mode (dia-PASEF). While the matched, confounder-controlled principle should generalize, the specific magnitudes of the depth advantage, CV differential, and divergence-zone counts reported here may vary across instruments, samples, and acquisition modes. Instruments of a different class, such as the Thermo Orbitrap and the Orbitrap Astral mass spectrometers, differ in their mass analyzer and acquisition characteristics. Whether the platform differences we report hold, narrow, or widen on those instruments is untested here and remains an open question that direct comparison on matched samples would resolve. Second, the CV comparison is computed on protein groups shared between platforms, so it does not reflect either platform’s CV behavior on its own divergence-zone protein groups. Third, we did not test non-tryptic enzymes, PTM-heavy datasets, or open-search configurations, which may produce different platform behavior. Fourth, our results are specific to SN-21 with the Kuiper search engine; earlier versions running the Pulsar engine may not reproduce the same behavior. Finally, the matched baseline isolates the search-engine and quantifier contribution by design, so the depth figures reported here characterize that isolated contribution rather than platform performance when cross-run transfer features are active, as they are by default in SN-21 and when DIA-NN is run with Match-Between-Runs enabled.

### Recommendations for Academic and Industry Workflows

The concrete recommendations that follow depend on workflow priorities. For academic laboratories pursuing discovery-driven interactome or unbiased-proteome characterization, we recommend running both platforms, treating the intersection as the high-confidence set, and treating each platform’s divergence zone as a hypothesis-generation set to be interpreted with appropriate skepticism informed by mechanistic characterization (see Figure 6). The specific pattern we observed, in which SN-only proteins live at the low-abundance edge of the dynamic range with lower detection completeness, applies here and can be applied as an internal QC filter. For industrial laboratories where low false discovery rate is paramount, and downstream statistical inference must be defensible, we recommend FP-24.0 for its 4.1 to 6.4 percentage-point tighter median CVs across all three fractions, and its explicit visibility into rescoring features via MSBooster. When using SN-21 in an industrial setting, we recommend running it in directDIA+ (Deep) mode with a fraction-restricted hybrid library and cross-run transfer configured to match the comparator platform. Regardless of platform choice, we recommend that head-to-head benchmark work explicitly report the analysis mode used on each platform and disable cross-run transfer features symmetrically to isolate the search-engine and quantifier contribution from the transfer-feature contribution.

## Abbreviations

AR: androgen receptor
CV: coefficient of variation
DDA: data-dependent acquisition
DIA: data-independent acquisition
dia-PASEF: data-independent acquisition parallel accumulation serial fragmentation
FDR: false discovery rate
FP: FragPipe 24.0
GO: Gene Ontology
iRT: indexed retention time
LC-MS/MS: liquid chromatography tandem mass spectrometry
MBR: Match-Between-Runs
PG: protein group
PSM: peptide-spectrum match
SN: Spectronaut 21.

## ASSOCIATED CONTENT

### Supporting Information

The Supporting Information is available free of charge at https://pubs.acs.org/doi/10.1021/acs.jproteome.XXXXXXX.

Supplementary Figures S1 to S3 with legends and the legends for Supplementary Tables S1 to S11 (PDF)

Supplementary Tables S1 to S11 (XLSX)

### Data Availability Statement

Raw dia-PASEF .d files and the processed outputs of both pipelines have been deposited to the ProteomeXchange Consortium via the PRIDE partner repository (45) with the dataset identifier PXD081449. The deposit includes the FragPipe 24.0 outputs, namely the diaTracer, DIA-NN, and report.pg_matrix.tsv result files, and the Spectronaut 21 outputs, namely the Pulsar-derived per-fraction session reports with their configuration files and the matched-condition and Match-Between-Runs protein-group matrices for the cytosolic, microsomal, and nuclear fractions. The project supplementary materials include analysis scripts, figure-generation scripts, and derived intermediate files.

## AUTHOR INFORMATION

### Authors

Lilliana Radoshevich. Department of Immunology and Genomic Medicine, National Jewish Health, Denver, CO, USA.

### Author Contributions

L.R. contributed to resources, methodology, investigation, and writing (review and editing). M.E.W. contributed to conceptualization, methodology, investigation, resources, writing (original draft and review and editing), supervision, and funding acquisition.

### Funding

This work was supported by R01GM143399 to M.E.W. The funders had no role in study design, data collection, analysis, interpretation, or the decision to submit the manuscript.

### Notes

M.E.W. received a temporary license to Spectronaut 21 from Biognosys to enable this study, and Biognosys reviewed and advised on the Spectronaut 21 analysis configuration. Biognosys had no role in data collection, data analysis, interpretation of the results, or the decision to submit the manuscript, and did not influence the study’s findings. The authors otherwise declare no conflict of interest.

## ACKNOWLEDGMENTS

We thank Alexey I. Nesvizhskii and Fengchao Yu of the University of Michigan for reviewing the FragPipe and MSFragger analysis configuration, and Monika Puchalska of Biognosys for reviewing the Spectronaut 21 analysis configuration. This independent review of each platform’s settings helped ensure that both pipelines were configured according to their developers’ recommendations. The authors thank Jimmy K. Eng (University of Washington Proteomics Resource) for reading the manuscript and for helpful discussions and thank Biognosys for access to a temporary license of the Spectronaut 21 search platform and for reviewing the Spectronaut 21 analysis configuration used in this study. The authors thank the g:Profiler team at the University of Tartu for the enrichment REST API. NCBI PubMed E-utilities provided literature-validation resources.

## Declaration of generative AI and AI-assisted technologies in the manuscript preparation process

During the preparation of this work, the authors used Claude (Anthropic) to assist with analysis scripting, figure generation from source data, reference verification, and manuscript editing. After using this tool, the authors reviewed and edited the content as needed and take full responsibility for the published article.

## Notes

### Competing Interest Statement

The authors have declared no competing interest.

