## Supplementary Figures for "Search-software choice shapes the discoverable androgen receptor proximal interactome in DIA proteomics"

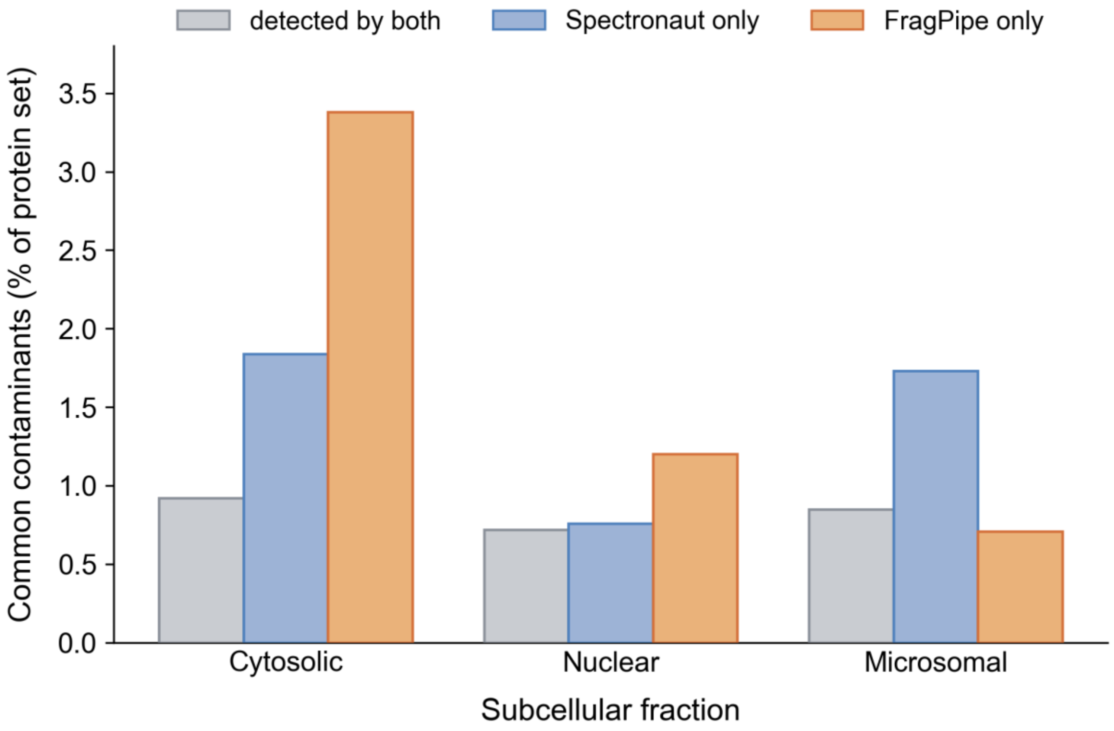


**Supplementary Figure S1.** Common-contaminant audit of the divergence-zone protein sets. Percentage of each protein set matching the universal protein contaminant library of Frankenfield et al. 2022, for proteins detected by both platforms, by SN-21 only, and by FP-24.0 only, in the cytosolic, nuclear, and microsomal fractions. The SN-only and FP-only rates remain low in every fraction and near the shared-proteome baseline, apart from a modestly higher cytosolic FP-only rate (4.23%, driven by keratin and immunoglobulin), confirming that neither divergence zone is meaningfully enriched for common contaminants relative to the shared proteome. The rates are reported in the Results.

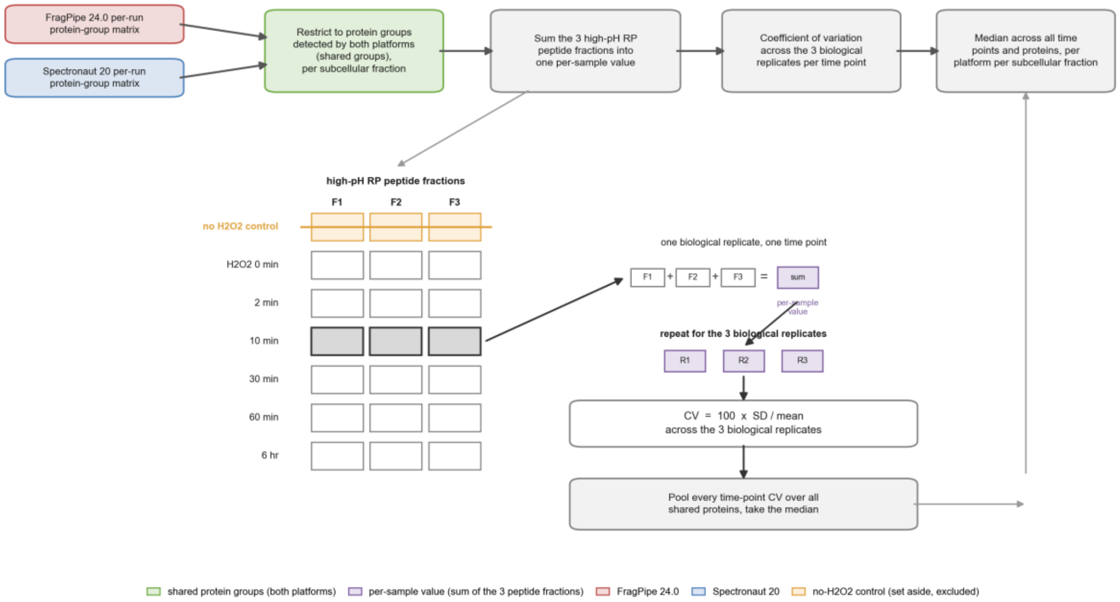


**Supplementary Figure S2.** Derivation of the biological-replicate coefficient of variation. Schematic of how the median coefficient of variation reported in Figure 4 is computed. Within each subcellular fraction, samples comprise three biological replicates, seven proximity-labeling time points, and three high-pH reversed-phase peptide fractions. The three peptide fractions of each sample are summed into a single per-sample value, after which the coefficient of variation is computed across the three biological replicates within each hydrogen-peroxide-labeled time point, on protein groups detected by both platforms, and pooled to a median per platform per subcellular fraction. The no-hydrogen-peroxide negative controls are set aside. Because MaxLFQ intensities are computed per run, the fraction summation approximates a sample-level quantity. The values are reported in the Results.

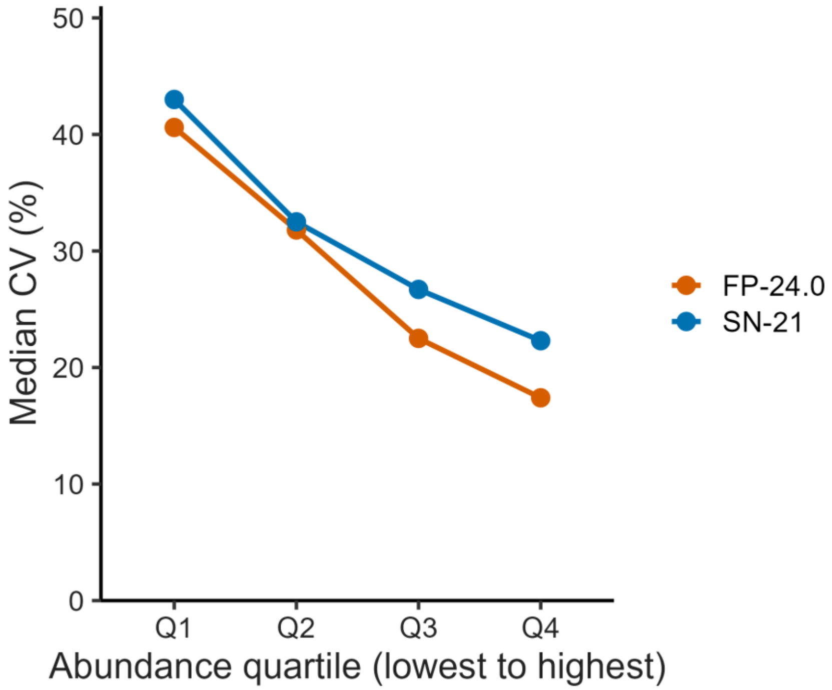
50
40
30
20
10
0
Median CV (%)
Q1
Q2
Q3
Q4
Abundance quartile (lowest to highest)
FP-24.0
SN-21

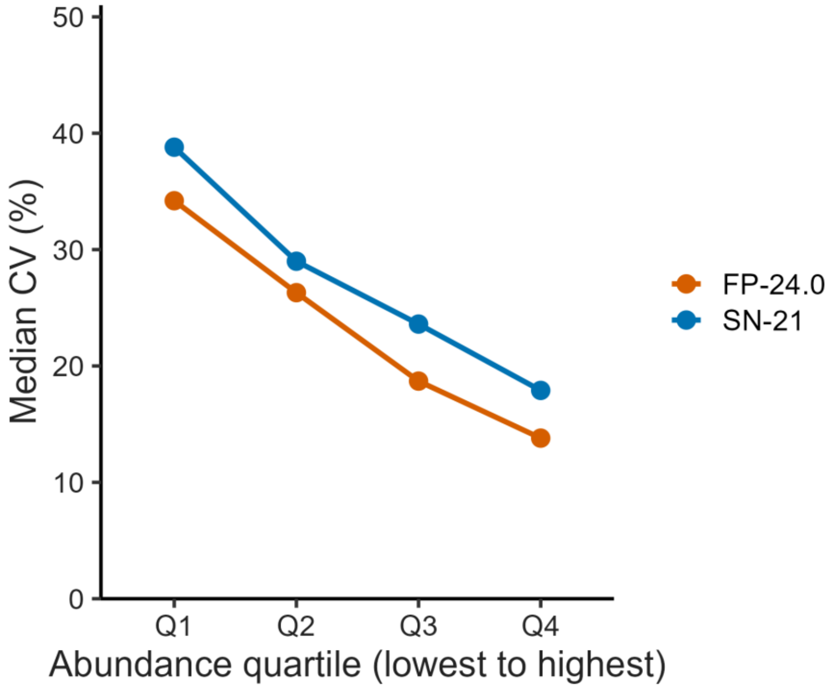
50
40
30
20
10
0
Median CV (%)
Q1
Q2
Q3
Q4
Abundance quartile (lowest to highest)
FP-24.0
SN-21

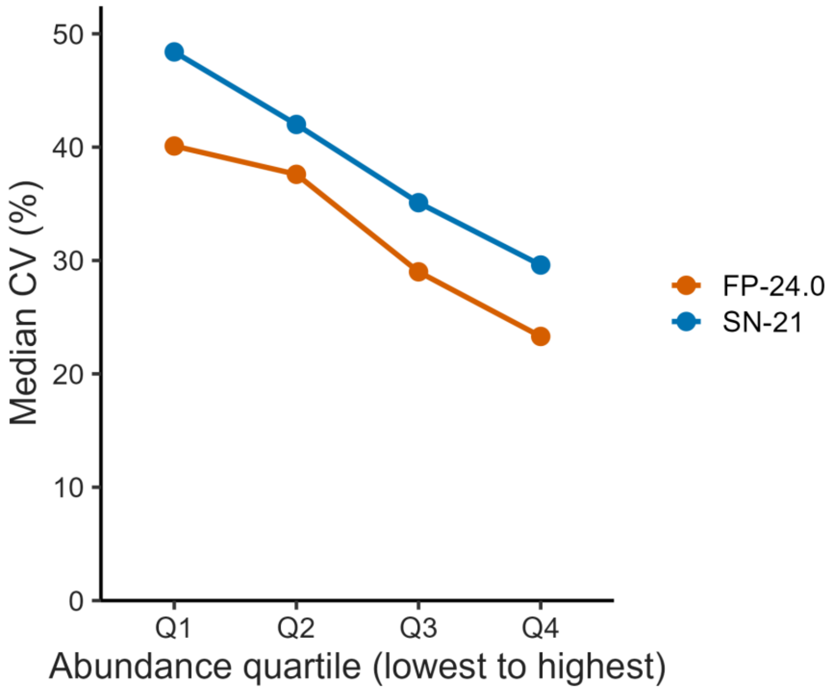


**Supplementary Figure S3.** Biological-replicate CV versus abundance, by platform. Median CV (%) of shared protein groups in each abundance quartile (lowest to highest, by each engine's own intensity) for FP-24.0 (orange) and SN-21 (blue), one panel per fraction. Computed by the Figure 4 method on abundance-stratified shared proteins (Supplementary Table S11). FP-24.0 is more precise than SN-21 in every quartile of every fraction, and the gap does not narrow toward high abundance.

### Supplementary Tables

Supplementary Tables S1 through S11 are provided as separate Microsoft Excel files.

**Supplementary Table S1.** Spectral library composition per fraction. DDA and DIA file counts, breakdown by biological replicate and sub-fraction.

**Supplementary Table S2.** FP-24.0 and SN-21 parameter map. Full side-by-side parameter mapping across all pipeline stages.

**Supplementary Table S3.** Per-fraction protein-group count and completeness summary. Referenced in the Results.

**Supplementary Table S4.** Full g:Profiler enrichment tables per fraction and per set, with the per-fraction set sizes and the AR-interactome subset analysis. Referenced in the Results.

**Supplementary Table S5.** Cross-fraction concordance analysis. Cross-fraction GO-term intersection and g:Profiler enrichment on the 146 SN-only and 10 FP-only genes recurring across all three fractions. Referenced in the Results.

**Supplementary Table S6.** Recovery of the known AR-interactome. Per-fraction recovery counts, the members recovered uniquely by each platform with their PubMed AR co-citation counts, and the shared members. Referenced in the Results.

**Supplementary Table S7.** Recurrent divergence-zone gene lists. The 146 SN-only and 10 FP-only genes recurring across all three fractions. Referenced in the Results.

**Supplementary Table S8.** Software versions and analysis scripts. Referenced in Data Availability.

**Supplementary Table S9.** Platform-unique AR-interactome members, network and enrichment. Gene lists recovered uniquely by each platform, their PubMed AR co-citation counts, STRING v11.5 physical member-to-member edges at score 0.70 or above, and g:Profiler enrichment against the measured proteome background. Referenced in the Results and Experimental Section.

**Supplementary Table S10.** Proteotypic peptide evidence for the FragPipe-only cytoskeletal proteins. Per-gene and per-fraction counts of uniquely-mapping (proteotypic) versus shared-only precursors for the tissue-restricted cytoskeletal family members in the cytosolic and nuclear FragPipe-only sets, together with the individual proteotypic peptide sequences, charges, and run-level detection so that the uniqueness of each can be verified independently. Referenced in the Results and Experimental Section.

**Supplementary Table S11.** Biological-replicate CV by abundance quartile. Median CV of shared protein groups, overall and in each abundance quartile, for FP-24.0 and SN-21 in all three fractions, computed by the Figure 4 method on abundance-stratified shared proteins. Referenced in the Results.
